# Pharmacological inhibition of lysine-specific demethylase 1 (LSD1) induces global transcriptional deregulation and ultrastructural alterations that impair viability in *Schistosoma mansoni*

**DOI:** 10.1101/829549

**Authors:** Vitor Coutinho Carneiro, Isabel Caetano de Abreu da Silva, Murilo Sena Amaral, Adriana S.A. Pereira, Gilbert O. Silveira, David da Silva Pires, Sérgio Verjovski-Almeida, Frank J. Dekker, Dante Rotili, Antonello Mai, Eduardo José Lopes Torres, Dina Robaa, Wolfgang Sippl, Raymond J. Pierce, M. Teresa Borrello, A. Ganesan, Julien Lancelot, Silvana Thiengo, Monica Ammon Fernandez, Amanda Roberta Revoredo Vicentino, Marina Moraes Mourão, Fernanda Sales Coelho, Marcelo Rosado Fantappié

**Author notes:** Epigenetic therapy: targeting histone de-methylation to control schistosomiasis. Division of Epigenetics, German Cancer Research Center, Im Neuenheimer Feld 580, 69120 Heidelberg, Germany (Vitor Coutinho Carneiro). Corresponding author: M.R. Fantappié, Instituto de Bioquímica Médica, Universidade Federal do Rio de Janeiro, CCS, Ilha do Fundão, Rio de Janeiro, 21941-902, Brazil.

## Abstract

Treatment and control of schistosomiasis still rely on only one effective drug, praziquantel (PZQ), and due to mass treatment, the increasing risk of selecting for schistosome strains that are resistant to PZQ has alerted investigators to the urgent need to develop novel therapeutic strategies. The histone-modifying enzymes (HMEs) represent promising targets for the development of epigenetic drugs against *Schistosoma mansoni*. In the present study, we targeted the *S. mansoni* lysine-specific demethylase 1 (SmLSD1), a transcriptional corepressor, using a novel and selective synthetic inhibitor, MC3935. We synthesized a novel and potent LSD1 inhibitor, MC3935, which was used to treat schistosomula or adult worms *in vitro*. By using cell viability assays and optical and electron microscopy, we showed that treatment with MC3935 affected parasite motility, egg-laying, tegument, and cellular organelle structures, culminating in the death of schistosomula and adult worms. *In silico* molecular modeling and docking analysis suggested that MC3935 binds to the catalytic pocket of SmLSD1. Western blot analysis revealed that MC3935 inhibited SmLSD1 demethylation activity of H3K4me1/2. Knockdown of SmLSD1 by RNAi recapitulated MC3935 phenotypes in adult worms. RNA-seq analysis of MC3935-treated parasites revealed significant differences in gene expression related to critical biological processes. Collectively, our findings show that SmLSD1 is a promising drug target for the treatment of schistosomiasis and strongly support the further development and *in vivo* testing of selective schistosome LSD1 inhibitors.

**Author Summary:** Schistosomiasis mansoni is a chronic and debilitating tropical disease caused by the helminth *Schistosoma mansoni*. The control and treatment of the disease rely almost exclusively on praziquantel (PZQ). Thus, there is an urgent need to search for promising protein targets to develop new drugs. Drugs that inhibit enzymes that modify the chromatin structure have been developed for a number of diseases. We and others have shown that *S. mansoni* epigenetic enzymes are also potential therapeutic targets. Here we evaluated the potential of the *S. mansoni* histone demethylase LSD1 (SmLSD1) as a drug target. We reported the synthesis of a novel and potent LSD1 inhibitor, MC3935, and show that it selectively inhibited the enzymatic activity of SmLSD1. Treatment of juvenile or adult worms with MC3935 caused severe damage to the tegument of the parasites and compromised egg production. Importantly, MC3935 proved to be highly toxic to *S. mansoni,* culminating in the death of juvenile or adult worms within 96 h. Transcriptomic analysis of MC3935-treated parasites revealed changes in the gene expression of hundreds of genes involved in key biological processes. Importantly, SmLSD1 contains unique sequences within its polypeptide chain that could be explored to develop a *S. mansoni* selective drug.

## Introduction

Schistosomes are large metazoan pathogens that parasitize over 200 million people worldwide, resulting in up to 300,000 deaths per year (1, 2). No efficacious vaccine is available for human schistosomiasis, and the control and treatment of the disease rely almost exclusively on praziquantel (PZQ), the only effective drug against schistosome species. Despite its efficacy, PZQ does not kill juvenile parasites, allowing reinfection (3), and there is a constant concern with the appearance of PZQ-resistant strains of *Schistosoma* (4–6). Thus, there is an urgent need to search for promising protein targets to develop new drugs.

Transcription factors and chromatin modifiers play primary roles in the programming and reprogramming of cellular states during development and differentiation, as well as in maintaining the correct cellular transcriptional profile (7). Indeed, a plethora of groundbreaking studies has demonstrated the importance of posttranslational modifications of histones for transcription control and normal cell development. Therefore, the deregulation of epigenetic control is a common feature of a number of diseases, including cancer (7).

The complexity of schistosome development and differentiation implies tight control of gene expression at all stages of the life cycle and that epigenetic mechanisms are likely to play key roles in these processes. In recent years, targeting the *Schistosoma mansoni* epigenome has emerged as a new and promising strategy to control schistosomiasis. The study of histone acetylation in *S. mansoni* biology and the effect of inhibitors of histone deacetylases (HDACs and SIRTs) or histone acetyltransferases (HATs) on parasite development and survival have demonstrated the importance of these enzymes as potential therapeutic targets (8–12).

Unlike histone lysine acetylation, which is generally coupled to gene activation, histone lysine methylation can have different biological associations depending on the position of the lysine residue and the degree of methylation (13). Patterns of specific lysine methyl modifications are achieved by a precise lysine methylation system, consisting of proteins that add, remove and recognize the specific lysine methyl marks. Importantly, histone lysine methylation (14–16) and demethylation (17) have been recently demonstrated to be potential drug targets against *S. mansoni*.

Lysine-specific demethylase 1 (LSD1) was the first protein reported to exhibit histone demethylase activity and has since been shown to have multiple essential roles in metazoan biology (18). LSD1 enzymes are characterized by the presence of an amine oxidase (AO)-like domain, which is dependent on its cofactor flavin-adenine dinucleotide (FAD), a SWIRM domain, which is unique to chromatin-associated proteins (19) and an additional coiled-coil TOWER domain (20). LSD1 is a component of the CoREST transcriptional corepressor complex that also contains CoREST, CtBP, HDAC1 and HDAC2. As part of this complex, LSD1 demethylates mono-methyl and di-methyl histone H3 at Lys4 (H3K4me1/2), but not H3K4me3 (21). In addition, when recruited by androgen or estrogen receptor, LSD1 functions as a H3K9 demethylase. Given the high level of expression and enzymatic activity of LSD1 in many types of tumors, there has been significant recent interest in the development of pharmacological inhibitors (22).

In our continuing effort to study the biology and therapeutic potential of epigenetic regulators in *S. mansoni*, we have found schistosome LSD1 (SmLSD1, Sm_150560) as a potential drug target (this paper). During the course of our investigation, a recent publication (17) described the repurposing of some anthracyclines as anti-Schistosoma agents, suggesting by *in silico* docking SmLSD1 as a putative target without any evidence of enzyme inhibition.

In the present work we show the SmLSD1-inhibitory activity of a novel synthetic LSD1 inhibitor MC3935 (100-fold more potent than the canonical human LSD1 inhibitor tranylcypromine, TCP). In addition, we show that the LSD1 inhibitor MC3935 was able to kill both adult worms and schistosomula *in vitro*. Importantly, silencing of SmLSD1 by dsRNAi partially recapitulated MC3935-treatment phenotypes in adult worms. Our RNA-seq analysis revealed a large-scale transcriptional deregulation in parasites that were treated with sublethal doses of MC3935, which could be the primary cause of the ultrastructure defects and death of *S. mansoni*. Together, these findings elucidate the biological relevance of histone lysine methylation in *S. mansoni* and provide insights into the therapeutic potential of SmLSD1 to control schistosomiasis.

## Materials and methods

### Ethics statement

Animals were handled in strict accordance with good animal practice as defined by the Animal Use Ethics Committee of UFRJ (Universidade Federal do Rio de Janeiro). This protocol was registered at the National Council for Animal Experimentation (CONCEA-01200.001568/2013-87) with approval number 086/14. The study adhered to the institution’s guidelines for animal husbandry.

### Protein alignment and phylogenetic relationships

Multiple-sequence alignment of the full-length proteins was performed using representatives of *C. elegans* (NP_493366.1), *D. melanogaster* (NP_649194.1), *H. sapiens* (NP_055828.2), *M. musculus* (NP_598633.2), *D. rerio* (XP_005158840.1)*, A. thaliana* (NP_187981.1), *S. japonicum* (TNN15244.1), *S. haematobium* (XP_012793780.1), and *S. mansoni* (Smp_150560) LSD1 protein sequences, as previously published (23). Pairwise comparisons to the reference followed by calculation of the maximum distance matrix resulted in an unrooted phylogenetic tree, which was visualized using Tree of Life v1.0 (24).

### Chemistry

Compound MC3935 (Supplementary Fig S1A) was synthesized by coupling of the racemic *tert*-butyl (2-(4-aminophenyl)cyclopropyl)carbamate, prepared as previously reported (25), with the commercially available 4-ethynylbenzoic acid, followed by acidic deprotection of the Boc-protected amine. ^1^H-NMR spectra were recorded at 400 MHz using a Bruker AC 400 spectrometer; chemical shifts are reported in ppm units relative to the internal reference tetramethylsilane (Me_4_Si). Mass spectra were recorded on an API-TOF Mariner by Perspective Biosystem (Stratford, Texas, USA). Samples were injected by a Harvard pump at a flow rate of 5-10 µL/min and infused into the electrospray system. All compounds were routinely checked by TLC and ^1^H-NMR. TLC was performed on aluminum-backed silica gel plates (Merck DC, Alufolien Kieselgel 60 F_254_) with spots visualized by UV light or using a KMnO_4_ alkaline solution. All solvents were reagent grade and, when necessary, were purified and dried by standard methods. The concentration of solutions after reactions and extractions involved the use of a rotary evaporator operating at a reduced pressure of ∼ 20 Torr. Organic solutions were dried over anhydrous sodium sulfate. Elemental analysis was used to determine the purity of the final compound 1 (MC3935) which was >95%. Analytical results were within 0.40% of the theoretical values. As a rule, the sample prepared for physical and biological studies was dried in high vacuum over P_2_O_5_ for 20 h at temperatures ranging from 25 to 40 °C. Abbreviations are defined as follows: dimethylformamide (DMF), *N*-(3-dimethylaminopropyl)-*N*′-ethylcarbodiimide (EDCI), 1-hydroxybenzotriazole hydrate (HOBt), triethylamine (TEA), ethyl acetate (EtOAc) and tetrahydrofuran (THF).

### Preparation of tert-Butyl-(trans-2(4-(4-ethynylbenzamido)phenyl)cyclopropyl) carbamate (3)

4-Ethynylbenzoic acid (135.4 mg, 0.93 mmol, 1.15 eq), EDCI (216.2 mg, 1.13 mmol, 1.4 eq), HOBt (152.4 mg, 1.13 mmol, 1.4 eq) and TEA (0.43 mL, 3.06 mmol, 3.8 eq) were added sequentially to a solution of 2 (200 mg, 0.805 mmol, 1.0 eq) in dry DMF (4.5 mL) (Fig. S1). The resulting mixture was then stirred for approximately 7 h at room temperature and, after completion of the reaction, quenched with NaHCO_3_ saturated solution (40 mL). The aqueous solution was extracted with EtOAc (4 x 25 mL); washed with 0.1 N KHSO_4_ solution (2 x 10 mL), NaHCO_3_ saturated solution (3 x 10 mL) and brine (3 x 5 mL); dried over anhydrous Na_2_SO_4_ and finally concentrated under vacuum. The crude product was then purified by column chromatography on silica gel eluting with a mixture EtOAc:hexane 25:75 to afford 3 as a pink solid (193 mg, 64%). ^1^H-NMR (400 MHz, CDCl_3_): δ 1.06-1.10 (m, 2H, - C*H_2_*-), 1.39 (s, 9H, -COO(C*H_3_*)*_3_*), 1.94-1.98 (m, 1H, Ar-C*H*-), 2.63 (m, 1H, -C*H*-NH-COO(CH_3_)_3_), 3.16 (s, 1H, *H*C≡C-), 4.78 (s, 1H, -N*H*-COO(CH_3_)_3_), 7.07-7.09 (d, 2H, *H*-Ar), 7.44-7.47 (d, 2H, *H*-Ar), 7.52-7.54 (d, 2H, *H*-Ar), 7.69 (s, 1H, Ar-CO-N*H*-Ar), 7.74-7.76 (d, 2H, *H*-Ar). MS (ESI), m/z: 377 [M + H]^+^.

### Preparation of N-(4-(trans-2-aminocyclopropyl)phenyl)-4-ethynylbenzamide hydrochloride (1, MC3935)

To a solution of 3 (125 mg, 0.332 mmol, 1 eq.) in dry THF (9 mL) was added 4N HCl in dioxane (5.4 mL, 21.6 mmol, 65 eq.) while cooling at 0 °C. Then, the resulting suspension was stirred at room temperature for approximately 1 h. Finally, the suspension was filtered off and washed over the filter in sequence with dry THF (1 x 3 mL) and dry diethyl ether (4 x 3 mL) to afford 1 (MC3935) as a slightly pink solid (90.8 mg, 87.5%). ^1^H-NMR (400 MHz, DMSO-*d_6_*): δ 1.16-1.21 (m, 1H, -CH*H*-), 1.33-1.38 (m, 1H, -C*H*H-), 2.26-2.31 (m, 1H, -C*H*-Ar), 2.77-2.80 (m, 1H, -C*H*-NH_2_·HCl), 4.43 (s, 1H, *H*C≡C-), 7.14-7.16 (d, 2H, *H*-Ar), 7.63-7.65 (d, 2H, *H*-Ar), 7.70-7.72 (d, 2H, *H*-Ar), 7.95-7.98 (d, 2H, *H*-Ar), 8.28 (br s, 3H, -CH-N*H_2_·H*Cl), 10.33 (s, 1H, Ar-CON*H*-Ar). MS (ESI), m/z: 277 [M + H]^+^. Anal. (C_18_H_17_ClN_2_O) Calcd. (%): C, 69.12; H, 5.48; Cl, 11.33; N, 8.96. Found (%) C, 69.26; H, 5.50; Cl, 11.27; N, 8.87.

### Biochemistry

Human LSD1 (KDM1A) (lysine (K)-specific demethylase 1A) was purchased from BPS Bioscience (catalog No 50097). Monomethylated histone peptide H3K4N(CH_3_) was purchased from Pepscan and horseradish peroxidase (HRP) from Pierce (Catalog No. 31490). The reagents for buffer preparation were purchased from Merck, Netherlands. Reactions were conducted in black 96-well flat-bottom microplates (Corning^®^ Costar^®^). The fluorescence measurements were carried out in a Synergy H1 Hybrid Multi-Mode Microplate Reader (BioTek, USA) and the gain setting of the instrument was adjusted to 70.. GraphPad Prism 5.0 was used for determination of the half-maximal inhibitory concentration (IC_50_). Nonlinear regression was used for data fitting.

### Human LSD1 inhibition assay

Compound 1 (MC3935) was screened for inhibition against human recombinant LSD1. The *in vitro* assay was based on the oxidative demethylation of the monomethylated histone peptide H3K4N(CH_3_) via a FAD/FADH_2_ mediated reduction of O_2_ to H_2_O_2_. The remaining LSD1 activity was monitored via the detection of the amount of H_2_O_2_ formed. This was done by horseradish peroxidase (HRP), which reduces H_2_O_2_ to H_2_O using Amplex Red^2^ as the electron donor. The resulting product, resorufin, was highly fluorescent at 590 nm. The inhibition assay was performed as described previously (26). The compound was preincubated at different concentrations with LSD1 for 15 min at room temperature in the presence of HRP-Amplex Red. The substrate was then added, and the fluorescence was measured for 30 min.

### Homology modeling

The amino acid sequence of SmLSD1 was retrieved from Unipro (27) (accession number: G4VK09). Subsequently, alignment of SmLSD1 and HsLSD1 sequences was performed using MOE version 2018.01 (*Molecular Operating Environment (MOE)*, 2018.01; Chemical Computing Group Inc., Canada). Long inserts in the sequence of SmLSD1 (aa: 205-271, 324-385, 828-860, 883-910, and 965-983) were deleted and consequently not modeled. The saved alignment file was used to generate a homology model of SmLSD1 based on the cocrystal structure of hLSD1 with MC2580 (PDB ID 2XAS) (26) using MODELLER 9.11 (28).

### Ligand preparation

Similar to the observed adduct of the analogous tranylcypromine derivative MC2584 with FAD (PDB ID 2XAQ (29)), an N5 adduct of MC3935 with FAD was generated in MOE using only the flavin ring of FAD. The generated adduct was then cured using LigPrep (Schrödinger Release 2018-1): protonation states were assigned at pH 7±1 using Epik, tautomeric forms, as well as possible conformers were generated, and energy minimized using the OPLS03 force field. As a result, 25 low-energy conformers were generated using the bioactive search module implemented in Schrödinger.

### Protein preparation

The generated homology model of SmLSD1 was prepared with Schrödinger’s Protein Preparation Wizard (Schrödinger Release 2018-1); where hydrogen atoms were added and the hydrogen bond network was optimized. The protonation states at pH 7.0 were predicted using the PROPKA tool in Schrödinger, and the structure was subsequently subjected to a restrained energy minimization using the OPLS03 force field (RMSD of the atom displacement for terminating the minimization was 0.3 Å).

### Docking

The receptor grid preparation for the docking procedure was carried out by assigning the coordinates of the cut cocrystallized adduct (only the flavin ring was kept in FAD) as the centroid of the grid box. Molecular docking was performed using Glide (Schrödinger Release 2018-1) in the Standard Precision mode. A total of 20 poses per ligand conformer were included in the postdocking minimization step, and a maximum of one docking pose was stored for each conformer.

### Parasite stock

The Belo Horizonte strain of *Schistosoma mansoni* (Belo Horizonte, Brazil) was maintained in the snail (*Biomphalaria glabrata*) as the intermediate host and the golden hamster (*Mesocricetus auratus*) as the definitive host (30). Female hamsters aged 3–4 weeks, weighing 50–60 g, were infected by exposure to a *S*. *mansoni* cercarial suspension containing approximately 250 cercariae using intradermal injection. The adult worms were obtained by hepatoportal perfusion at 42–49 days postinfection. Cercariae were released from infected snails and mechanically transformed to obtain schistosomula *in vitro* (31).

### Treatment of *S. mansoni* with LSD1 inhibitors

Schistosomula or adult worms were treated with different concentrations of LSD1 inhibitors, as indicated in the figure legends. For each treatment condition, 10 worm pairs were maintained in 60-mm diameter culture dishes in 2 mL of culture medium (medium M169 (Gibco) supplemented with 10% fetal bovine serum (Vitrocell), penicillin/streptomycin, amphotericin and gentamicin (Vitrocell). Schistosomula were maintained in 96-well or 24-well culture plates, depending on the experiment, with 200 µL or 1 mL of culture medium M169 (Gibco), respectively, supplemented with 2% fetal bovine serum (Vitrocell), 1 µM serotonin, 0.5 µM hypoxanthine, 1 µM hydrocortisone, 0.2 µM triiodothyronine, penicillin/streptomycin, amphotericin and gentamicin (Vitrocell). Parasites were maintained at 37 °C in 5% CO_2_ with a humid atmosphere. The medium containing the LSD1 inhibitors or DMSO (vehicle) was refreshed every 24 h during the treatment period (1– 4 days).

### Viability assay

An inverted stereomicroscope (Leica M80) was used to evaluate the physiology and behavior of the parasites. Parasites were observed every 24 h, and representative images and videos were acquired. Schistosomula motility, light opacity, and membrane integrity were evaluated. Adult worm motility, pairing state, adherence to dish surface, and egg laying were monitored and determined. The viability was determined using the CellTiter-Glo Luminescent Cell Viability Assay (Promega) (32). Total cell lysates from one thousand schistosomula or 10 adult worms (paired or unpaired) were submitted to an ATP dosage. Eggs laid on the plates were quantified daily.

### SmLSD1 mRNA quantification

After treatment, the total RNA was extracted using a RiboPure kit (Ambion) followed by DNase treatment (Ambion) and cDNA synthesis (Superscript III, Invitrogen), following the manufacturer’s instructions. The resulting cDNA was diluted 10-fold in water and qPCR amplification was performed with 5 µL of diluted cDNA in a total volume of 15 µL using SYBR Green Master Mix (Life Technologies) and specific primer pairs (SmLSD1_qPCR_F: 5’ – CCACTTCAAACTGCCCTGTC – 3’ and SmLSD1_qPCR_R: 5’ – TCATCTTGATCCCAATGACGT – 3’, SmTubulin_qPCR_F: 5’ – GGATTTGACGGAATTCCAAA – 3’ and SmTubulin_qPCR_R: 3’ – AACGCTTAACTGCTCGTGGT – 3’) designed for *S*. *mansoni* genes by Primer3 online software (http://www.bioinformatics.nl/cgi-bin/primer3plus/primer3plus.cgi). QuantStudio 3 Real-Time PCR System (Applied Biosystems) was used. The results were analyzed by the comparative Ct method, and the statistical significance was calculated with the student t-test.

### Western blotting

Nuclear protein extracts from schistosomula or adult worms were prepared as previously described (31). From each sample, 10 µg of each extract was loaded on 7-12% precast SDS-polyacrylamide gels (Bio-Rad). After transference, membranes were blocked with Tris-buffered saline (TBS) containing 0.1% Tween 20 and 2% bovine serum albumin (TBST/2% BSA) and then probed overnight with specific antibodies in TBS/2% BSA. Membranes were washed with TBST and incubated for 1 h in TBST/2% BSA with secondary antibody (Immunopure goat anti-mouse # 31430, Thermo Scientific, and peroxidase-labeled affinity anti-rabbit # 04-15-06, KPL). After washing the membranes in TBST, the bands were visualized and images were recorded with the Amersham Imaging System (GE Healthcare), and quantified with ImageJ software (NIH). Histone monoclonal antibodies used were anti-H3K4me1 (#5326, Cell Signaling), anti-H3K4me2 (#9725, Cell Signaling) and anti-H3K4me3 (#9727, Cell Signaling), following the manufacturer’s instructions. For all antibodies, a 1:1000 dilution was used. For normalization of the signals across the samples, anti-histone H3 antibody (#14269, Cell Signaling) was used.

### Caspase 3/7 activity

The activity of caspase 3/7 was measured using the Caspase-Glo 3/7 assay kit (Promega) following the instructions. Cell lysates from schistosomula were obtained from 2,000 parasites cultivated in a 24-well plate with complete medium (as described above) and treated with MC3935 25 µM or vehicle (DMSO 0.25%). The luminescence was measured in a white-walled 96-well plate in a Wallac Victor2 1420 multilabel counter (PerkinElmer).

### TUNEL assay

Detection of DNA strand breaks in MC3935-treated schistosomula was performed using the *In situ* Cell Death Detection kit (Roche), as previously described (33). Schistosomula were fixed after 72 h treatment with MC3935 or DMSO. Parasites were mounted in a superfrost glass slide using Prolong with DAPI (Invitrogen) for nuclear visualization. Images were taken on a Zeiss Axio Observer Z1 (Zeiss) inverted microscope equipped with a 40X objective lens and an AxioCam MRm camera in ApoTome mode.

### Confocal laser scanning microscopy

For the confocal microscopy analysis, the adult worms were fixed and stained as previously described (34). Confocal scanning laser microscopy was performed on a Zeiss LSM 800 microscope equipped with a 488-nm HE/Ne laser and a 470-nm long-pass filter but without the reflection mode.

### Scanning and transmission electron microscopy

Scanning electron microscopy (SEM) and transmission electron microscopy (TEM) were performed to analyze ultrastructural alterations in the parasites. Adult worms or schistosomula were incubated with 25 µM MC3935 or 0.25% DMSO for 48 or 72h, and fixed, as previously described (34). For SEM analysis, the samples were dehydrated with increasing concentrations of ethanol and then dried with liquid CO_2_ in a critical-point dryer machine (Leica EM CPD030, Leica Microsystems, Illinois, USA) (35). Treated specimens were mounted on aluminum microscopy stubs and coated with gold particles using an ion-sputtering apparatus (Leica EM SCD050, Leica Microsystems). Specimens were then observed and photographed using an electron microscope (FEI QUANTA 250, Thermo Fisher Scientific). TEM analysis was performed on a Tecnai G2 microscope (FEI Company). Fixed specimens were washed in 0.1 M cacodylate buffer, pH 7.2; postfixed in 1% OsO_4_ and 0.8% K_3_Fe (CN)_6_; washed in 0.1 M cacodylate buffer, pH 7.2; dehydrated in a graded acetone series (20°–100° GL) for one hour each step and embedded in Polybed 812 epoxide resin. Thin-sections (60 nm) were collected on copper grids and stained for 30 minutes in 5% aqueous uranyl acetate and for 5 minutes in lead citrate.

### Double stranded RNA interference (RNAi)

The coding sequence of lysine-specific histone demethylase 1 (SmLSD1) (GenBank accession #: XM_018797592.1) was amplified by PCR using the oligonucleotides SmLSD1_F_1 (5’ –GTCGTCCCGTAACTCCAGTG – 3’) with SmLSD1_R_1 (5’ – AACAGGCAAGGTTTCGGACA – 3’) and SmLSD1_F_2 (5’ – TGTCACACGATGGAGAACTG – 3’) with SmLSD1_R_2 (5’ – GAAGTGTAGATTTGTCGATTGTGAA – 3’) with adult parasite cDNA synthesized using 5 ng of total RNA as template. These amplicons (SmLSD1_1 and SmLSD1_2) were used in a second PCR (nested PCR), diluted 1:500, with the oligonucleotides containing an upstream T7 tail sequence, respectively (SmLSD1_1 amplicon with SmLSD1_3 oligos and SmLSD1_2 amplicon with SmLSD1_4 oligos): SmLSD1_F_3 (5’ – GGGTAATACGACTCACTATAGGCCATCTCATACGTCGGTCCA – 3’) with SmLSD1_R_3 (5’ – GGGTAATACGACTCACTATAGGCTTTCAGCAGGCGTCAGAGTA– 3’) and SmLSD1_F_4 (5’ – GGGTAATACGACTCACTATAGGGACTCGTATGTTGCTGTCGGAG – 3’) with SmLSD1_R_4 (5’ – GGGTAATACGACTCACTATAGGCGGCTTCACGTAGACCACTT – 3’).

The *GFP* gene was used as a nonrelated dsRNA control and was amplified from pEGFP-N3 with the oligonucleotides: GFP_F_woT7 (5’ – AGCAGAGCTGGTTTAGTGAACC – 3’) with GFP_R_woT7 (5’ – TTATGATCTAGAGTCGCGGCCG – 3’). This amplicon was used in a nested PCR with the oligonucleotides containing a T7 tail: GFP_F_wT7 (5’ – GGGTAATACGACTCACTATAGGGGGATCCATCGCCACCATGGT – 3’) with GFP_R_wT7 (5’ – GGGTAATACGACTCACTATAGGGTTACTTGTACAGCTCGTCCATGCCG – 3’). Double-stranded RNA (dsRNA) was synthesized from templates of amplified PCR with oligonucleotides containing the T7 tail. The dsRNA was delivered by soaking the parasite couples in media containing 30 µg/mL of the desired dsRNA, and everyday, 70% of the medium was changed to a fresh medium also containing 30 µg/mL of dsRNA. At the end of the 2^nd^, 4^th^, and 7^th^ days of incubation, parasites were collected, washed twice in PBS and stored in RNAlater (Ambion) until RNA extraction. At the end of the 2^nd^, 4^th^ and 7^th^ days of incubation the total number of eggs, the number of parasites attached to the plate and the number of couples still paired were quantified. For the H3K4me1 and H3K4me2 western blotting, parasites were collected on the 7^th^ day of incubation with dsRNA and stored in PBS at -80 °C. The viability of the parasites was determined on the 7^th^ day of incubation as described above. Male and female adult worms were ground with glass beads in liquid nitrogen for 5 minutes. RNA extraction and cDNA synthesis were performed as described above. qPCR results were analyzed by the comparative Ct method. Real-time qPCR data were normalized in relation to the level of expression of the Smp_090920 (Fwd 5’ – CACCAGCTCATCATAAATAATCCA – 3’, Rev 5’ – TAGCATCCTGAAAGCCACGA – 3’) and Smp_062630 (Fwd 5’ – GGAATGATGTGGCCGATAGT – 3’, Rev 5’ – CGCAGAGATTGGCTAAATTG – 3’) reference genes.

### RNA-Seq data analysis

The general outline of the bioinformatics pipeline used for the analysis of the RNA-Seq data is completely described in Pereira et al. (36), including the three different statistical approaches that were used to obtain lists of differentially expressed genes, and considering as the final set only those genes that were listed at the intersection of the three sets. We used the same versions of genome and transcriptome annotation, including the use of a metagenes transcriptome to deal with isoforms, as previously described (36). All software parameters were as described (36) except for Trimmomatic (37) HEADCROP 12 and MAXINFO 60, since we decided to prioritize longer reads. Each replicate sample of adult worms has generated from 26 to 39 million paired-end 150-bp reads; for schistosomula, a total of 30 to 40 million reads was obtained per replicate sample.

## Results

### *Schistosoma mansoni* lysine specific-demethylase 1

The *Schistosoma mansoni* LSD1 (SmLSD1) contains all three canonical structural, and functional domains (SWIRM, amino-oxidase-like and TOWER domains) (Fig 1A) found in the LSD1 protein family. We examined in detail the protein alignment between SmLSD1 and human LSD1 (hLSD1) (Supplementary Fig S2), particularly since the latter is a well-defined drug target (18) and carried out a limited phylogenetic study including further orthologs. In this regard, our phylogenetic tree revealed that the SmLSD1 protein was closer to human LSD1 than to plant or nematode LSD1 (Fig 1B). It is worth noting that among the five LSD1 homologs tested only SmLSD1 presented unique sequences within all LSD1 functional domains (Fig 1A, purple segments and Supplementary Figure S2, dashed lines), which could be explored to develop a *S. mansoni*-selective drug.

**Figure 1.**
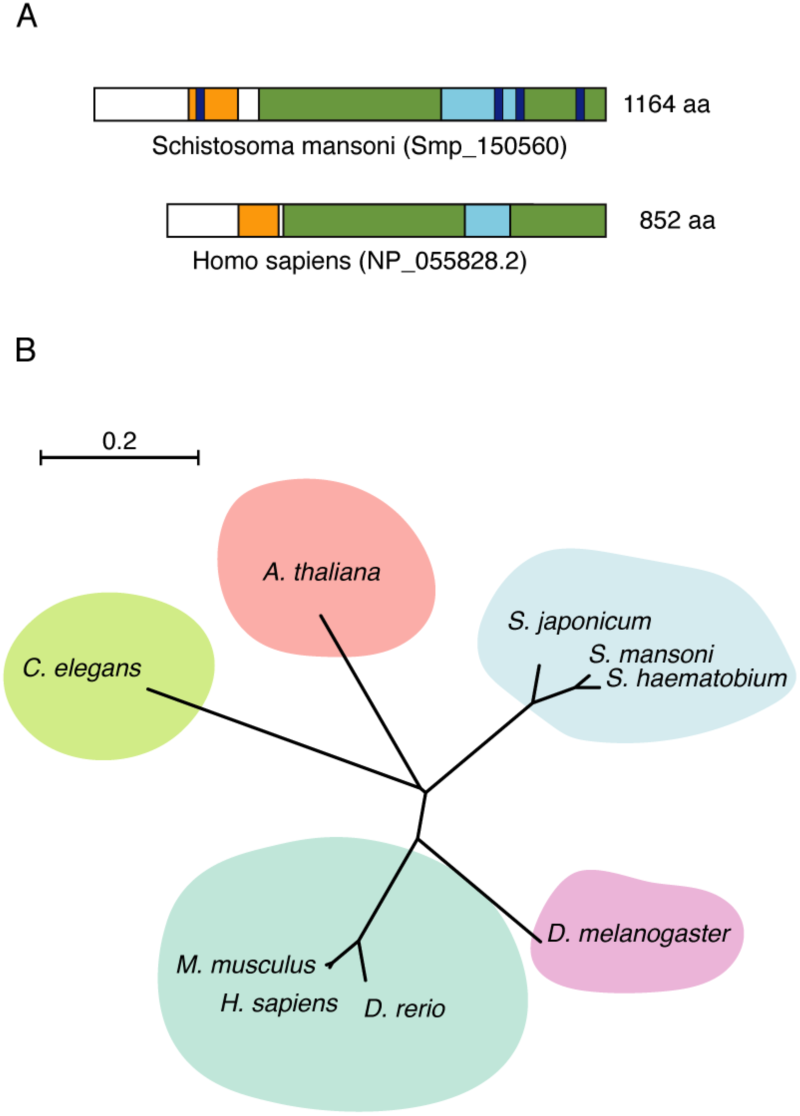
Overview of SmLSD1 protein domains and conservation. (A). Schematic representation of the full-length SmLSD1 protein (Smp_150560, top scheme), depicting the conserved functional domains: the SWIRM domain (orange), amine oxidase-like domain (green), the TOWER domain (blue), and schistosome unique sequences (purple). The full-length of the human LSD1 protein (bottom scheme) is also shown for comparison. (B). Unrooted phylogenetic tree representation was made using the ClustalW2 program and visualized with https://itol.embl.de/.https://itol.embl.de/. Tree scale: 0.2

### Viability of *Schistosoma mansoni* after MC3935 treatment

Schistosomula were obtained by mechanical transformation of cercariae, and adult worms were recovered by perfusion of infected hamsters (Fig 2A). We screened a series of LSD1 inhibitors (all these small compounds were synthesized based on the scalffold of tranylcypromine (TCP), a well-tested irreversible LSD1 inhibitor) to evaluate their schistosomicidal activity. Interestingly, all compounds at a final concentration of 25 µM showed toxicity against the juvenile form of schistosomula at 72 h (Supplementary Fig S3A) or adult worms at 96 h of cultivation (Supplementary Fig S3B). Interestingly, TCP showed the least toxic activity, whereas MC3935 was the most potent compound (Supplementary Fig S3A and B). Therefore, MC3935 was chosen for all further analyses in this study. We showed that MC3935 was able to inhibit the catalytic activity of the recombinant human LSD1 protein (Supplementary Fig S1B), revealing a 1,000 fold higher inhibitory activity than TCP, proving it as a *bona fide* LSD1 inhibitor. The toxic effect of MC3935 on schistosomula or adult worm pairs was further confirmed (Fig 2 B and C and Supplementary Fig S3C). A significant loss of viability at 10 µM and a complete loss of viability (what we judged death) at 25 µM MC3935 was observed in schistosomula or adult worms after cultivation for 72 or 96 h, respectively (Supplementary Fig S3C). These results were confirmed with videos (Supplementary videos S1-S4), which showed nearly 100% of the schistosomula had a complete lack of motility, high granularity and altered body shape (supplementary video S2) when compared to healthy schistosomula that were treated with DMSO only (supplementary video S1). MC3935-treated-adult worms also showed significant alterations when compared to the control (Fig 2D and Supplementary video S3), which included unpairing, lack of adherence, extremely low motility, vitellaria involution and no egg laying (Fig 2D and supplementary video S4). In order to evaluate whether the lack of eggs (Fig 2D; Egg laying) was exclusively due to the separation of the worms (Fig 2D; Pairing), we performed an additional experiment in which only adult worms that were kept coupled were maintained in culture, followed by egg counting. Worm pairs that were not treated, or treated with DMSO laid a significant number of eggs, while worm pairs that received the treatment of MC3935 laid no eggs whatsoever (Supplementary Fig S4A and B).

**Figure 2.**
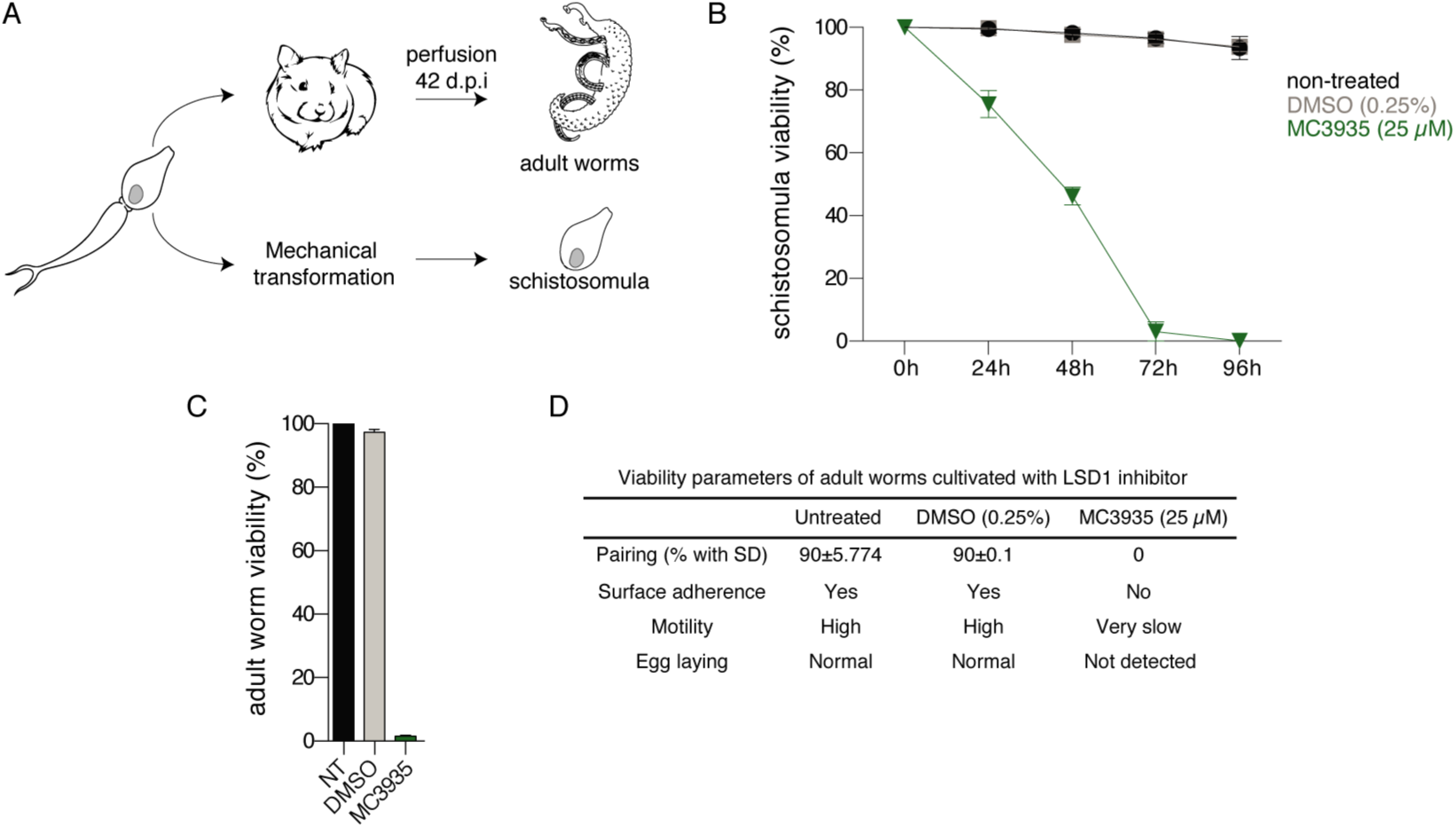
LSD1 inhibition is detrimental to *Schistosomam mansoni* survival. (A). Simplified scheme of the acquisition of the two developmental stages of the parasite used in this study. Cercariae were harvested from infected snails and used to either infect hamsters or mechanically transformed into schistosomula for *in vitro* culture. Hamsters were perfused 42 - 49 days postinfection to harvest adult worm pairs. (B). The relative ATP dosage (%) of schistosomula treated with 25 µM MC3935 (green line) was measured every 24 h (up to 96 h). Schistosomula given DMSO or nothing are shown in gray and black lines, respectively. (C) Relative ATP dosage (%) of adult worm pairs treated with 25 µM MC3935 for 96 h. NT, nontreated adult worm pairs. The results of three independent experiments are shown, and error bars represent the standard deviation (SD). (D) Evaluation of the viability of adult worm pairs treated with DMSO or 25 µM MC3935 for 96 h. Several parameters of adult worm viability were monitored daily until day 4, using an optical microscope equipped with a digital camera. Details for these classifications are described in the methods section. These viability parameters were reviewed and scored by two independent observers. Videos of the control or MC3935-treated worms to confirm the described scores are available (in Supplementary videos).

### Molecular docking and catalytic inhibition of SmLSD1 by MC3935

Since no crystal structure of SmLSD1 is yet available, a homology model of the parasite’s enzyme was first generated using an available crystal structure of the orthologous human LSD1. By analyzing the reported crystal structures of hLSD1 in complex with covalently bound tranylcypromine (TCP) derivatives, two crystal structures were found to be most relevant: a crystal structure of the N5 adduct of the highly analogous MC2584, which only lacked the ethynyl group found in MC3935, and the crystal structure with the N5 adduct of the bulky tranylcypromine derivative MC2580 (PDB IDs 2XAQ and 2XAS; respectively (29)). The latter crystal structure (PDB ID 2XAS) was preferred for the use as a template for the homology model since the conformation of residues Glu682 and Asp669 resulted in a more open binding site. Sequence alignment of SmLSD1 and hLSD1 showed an overall sequence identity of 44.1%, while the binding site of the FAD-ligand adduct shared an 80.4% sequence identity. In order to predict the binding mode of MC3935 to SmLSD1, the N5 adduct of this tranylcypromine derivative with the flavin ring of FAD was first generated similar to the N5 adduct of the analogous MC2584, and docking was subsequently performed into the homology model of SmLSD1. The obtained docking pose showed that the N5 adduct of MC3935 adopted a similar orientation in the binding site as observed with MC2584 (Fig 3A) with the ethynyl group embedded between Glu682 and Asp669. Notably, the binding site of SmLSD1 accommodating the tranylcypromine part of the adduct shared a 100% homology with the hLSD1 counterpart (Fig 3B).

**Figure 3.**
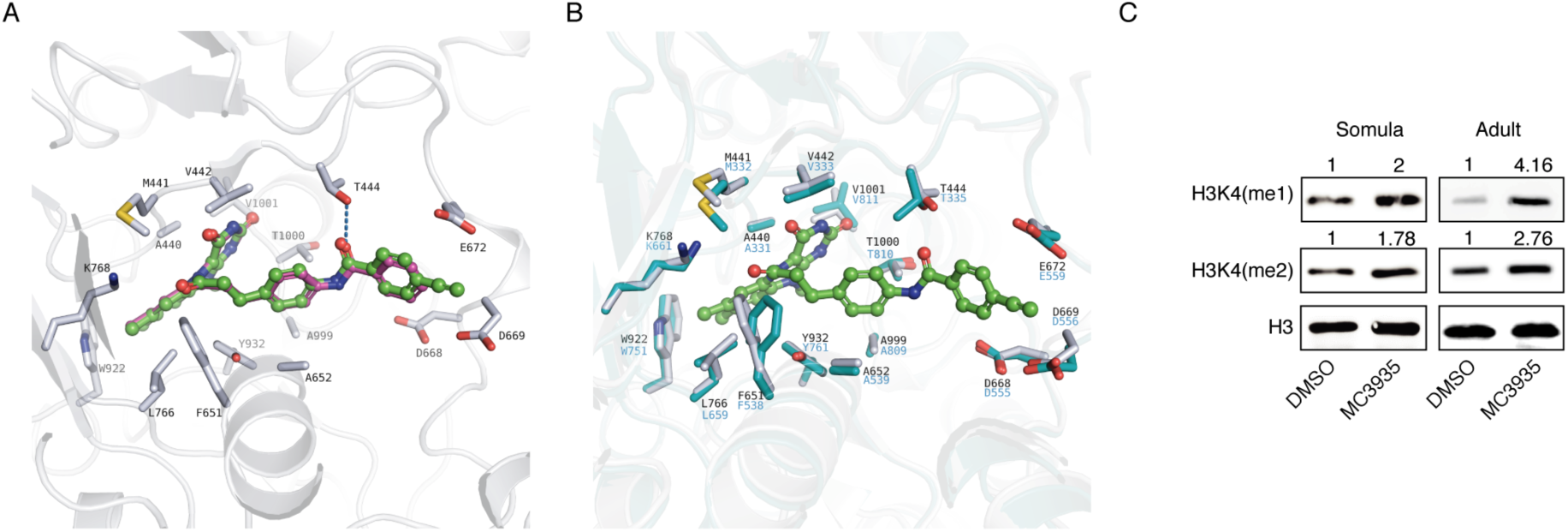
MC3935 binds to the catalytic pocket of SmLSD1 and inhibits its demethylase activity. (A). *In silico* molecular docking pose of the N5 adduct of MC3935 (in green) in the homology model of SmLSD1. The experimentally determined binding mode of the analogous MC2584 obtained by the superposition with the corresponding hLSD1 crystal structure (PD ID 2XAQ) is shown in purple. Only side chains of the SmLSD1 binding site are shown (white sticks). (B). Overlay of the SmLSD1 homology model (white sticks) showing the predicted binding mode of the MC3935 adduct with hLSD1 (cyan sticks; PDB ID 2XAS). (C) Western blot of total protein extracts from 72-hour-treated schistosomula (left panels) or 96 hour-treated adult worm pairs (right panels). Monoclonal antibodies against H3K4me1, H3K4me2 and H3 (as loading control) were used. Quantification of the bands (shown above each image) was done by densitometry (ImageJ, NIH software) normalized by the intensity of the H3 band. Western blots were performed from 5 independent biological replicates and one representative is shown here.

We next performed western blot analysis and showed that schistosomula or adult worms treated with MC3935 displayed higher band intensities of H3K4me1 or H3K4me2 marks when compared to the DMSO controls (Fig 3C), confirming the inhibition of SmLSD1 demethylase activity. Of note, the increase in H3K4me1 or H3K4me2 methylation was not due to the downregulation of SmLSD1 transcription (Supplementary Fig S5, qPCR graphs in A and B). Together, these data confirm that MC3935 inhibited SmLSD1 demethylase activity. Importantly, MC3935 treatments did not alter the H3K4me3 mark in schistosomula or adult worms (Supplementary Fig S5, western blots in A and B), pointing to a selective inhibition of LSD1-specific histone marks.

### Apoptosis in *S. mansoni* after SmLSD1 inhibition

The treatment of schistosomula with MC3935 significantly induced apoptosis, as detected by the 8-fold increase in the activities of caspases 3 and 7 (Fig 4A). In addition, the TUNEL assay indicated extensive double-strand DNA breaks (Fig 4B), as revealed by the green staining of the whole body of the worm treated with the inhibitor. Worms treated with DMSO showed no apoptotic activity, whatsoever (Fig 4A and B). These results are in agreement with our data from the ATP viability assay (Fig 2B, Supplementary Fig S3C) and our observations taken from the optical microscope (Supplementary videos S1 and S2).

**Figure 4.**
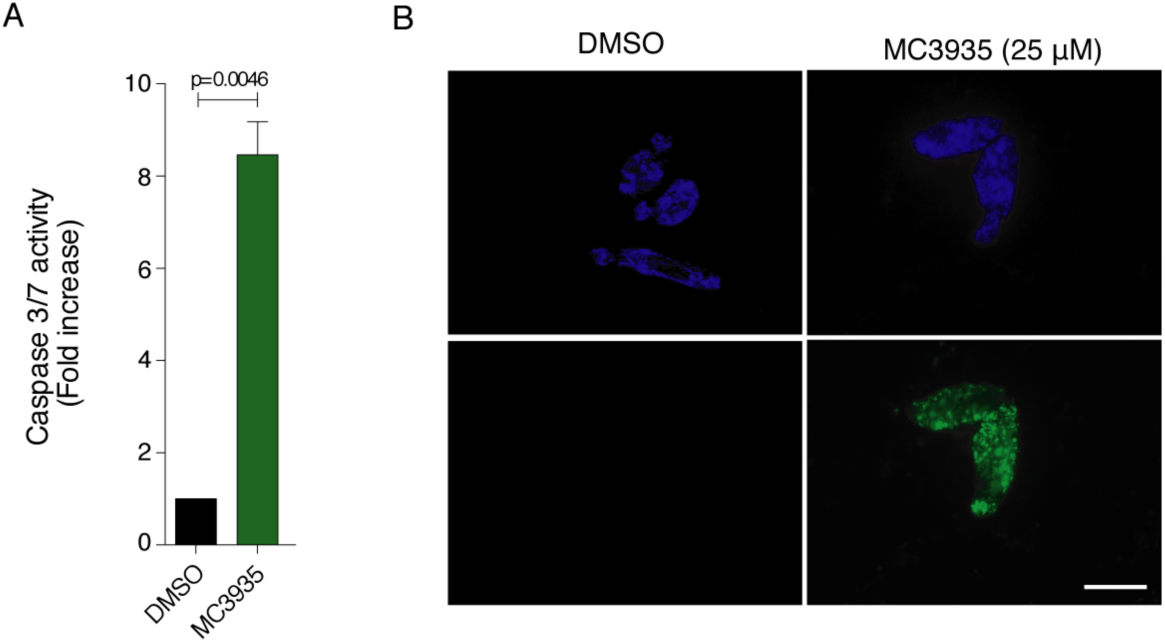
Inhibition of SmLSD1 triggers apoptosis. Twenty thousand schistosomula were cultivated in the presence of DMSO or 25 µM MC3935 for 72 h. (A) Caspase 3/7 activity was significantly increased in MC3935-treated parasites (green bar). (B) TUNEL assay of schistosomula incubated for 72 h with DMSO (left panels) or 25 µM MC3935 (right panels). Green parasites (lower panel) indicate double-strand DNA breaks. DAPI stains nuclear DNA, seen in blue (top panels). Scale bar: 50 µm.

### Tegumental damage and ultrastructural abnormalities of schistosomula after SmLSD1 inhibition

The results from our scanning electron microscopy clearly showed that inhibition of SmLSD1 by MC3935 induced severe erosions and fissures in the tegument of schistosomula (Fig 5B and D). Worms treated with DMSO revealed the typical healthy status of schistosomula (panel A), showing preserved tegumental spines (panel C). These images corroborate our conclusions that the MC3935 treatment led to schistosomula death. In this respect, it is reasonable to assume that the survival of these worms (note in panels B and D the depth of tegumental damage) could not be rescued even by the eventual withdrawal of the inhibitor.

**Figure 5.**
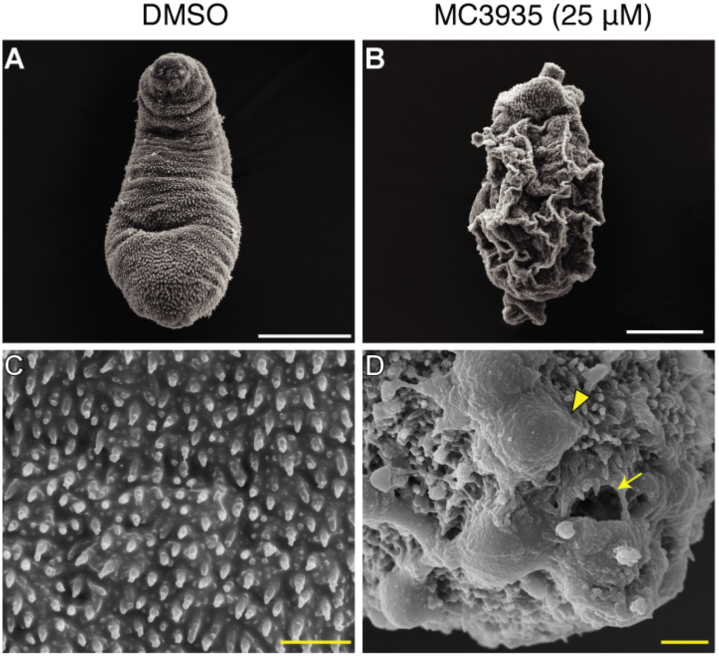
Inhibition of SmLSD1 leads to tegumental damage of schistosomula. Schistosomula were treated with 0.25% DMSO (left columns) or 25 µM MC3935 (right columns) for 48 h. Scanning electron microscopy (SEM) images of the tegument in lower (A and B) and higher magnification (C and D). Severe tegumental erosions (arrowhead), and fissures (arrow) are seen at higher magnification (D). Scale bars: white (5 µm) and yellow (1 µm).

Transmission electron microscopy of DMSO-treated schistosomula revealed preserved ultrastructures in the worms (Fig 6, panels A, C, E, G and I), such as tegumental spines (black arrows), outer tegument (*), tegument basal lamina (b), circular muscle (cm), longitudinal muscle (lm), mitochondria (m) and nuclei (n). In MC3935-treated schistosomula, extensive ultrastructural disorganization of the tegument was seen, such as a lack of the outer tegument (Fig 6, panel B, asterisks) and tegumental spines (panel B, black arrow). Significant loss of the muscle layers was also observed (compare lm and cm in panels A, C with panels B, D).

**Figure 6.**
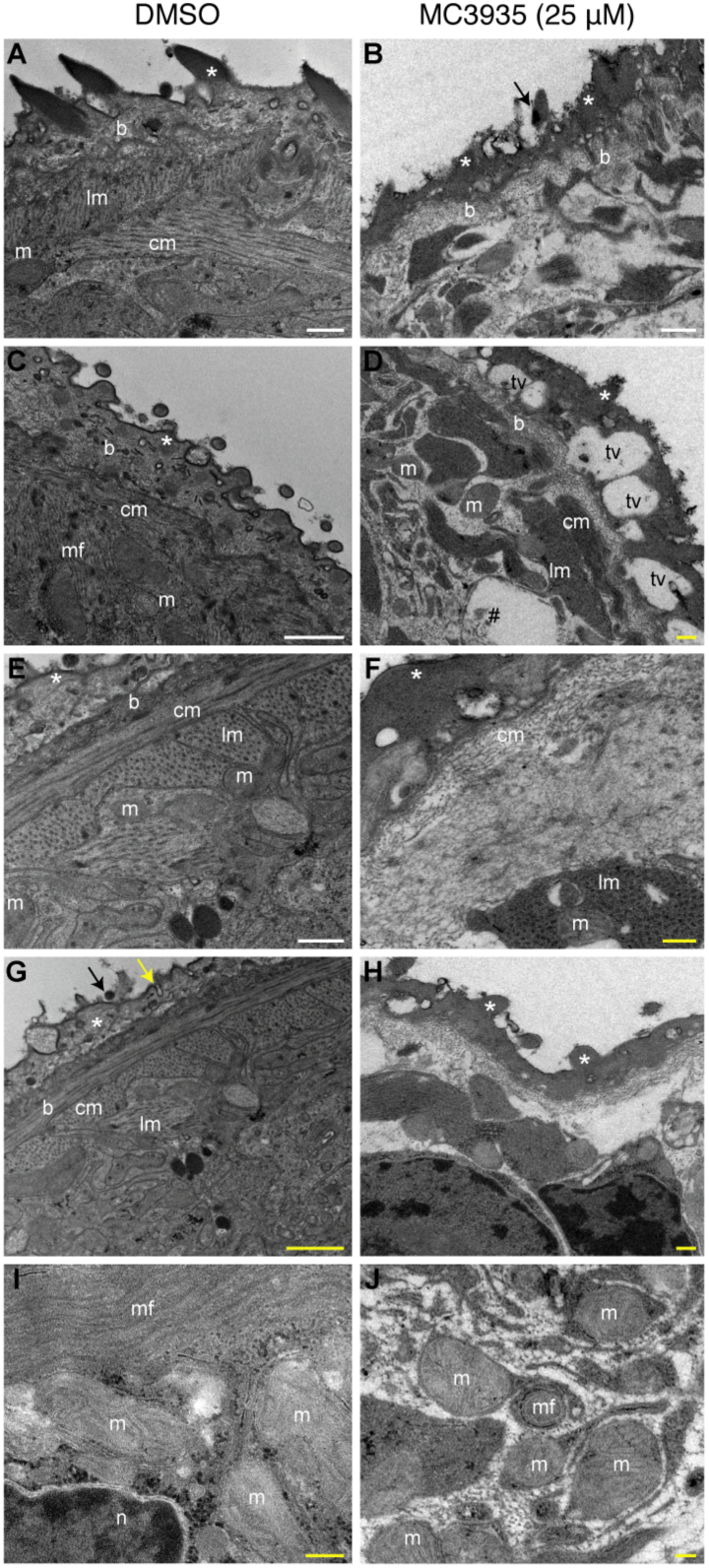
Inhibition of SmLSD1 leads to ultrastructural abnormalities in schistosomula. Transmission electron microscopy (TEM) of schistosomula treated with 0.25% DMSO (left column, panels A, C, E, G and I) or 25 µM MC3935 (right column, panels B, D, F, H and J) for 48 h. Symbols are as follows: tegumental spines (black arrows), outer tegument (*), tegument basal lamina (b), circular muscle (cm), longitudinal muscle (lm), mitochondria (m) and nucleus (n). In MC3935-treated schistosomula, an ultrastructural disorganization of the tegument is seen, lacking the outer tegument and tegumental spines (panel B, black arrow). A complete lack of the muscle layers (lm or cm) is also noted (panel B). Large vacuoles are observed in the more external region of the tegument of MC3935-treated parasites (panel D, tv). Internal vacuoles contain cellular debris (panel D, #). Significant thickening and higher electron density of the outer tegument is present, associated with the appearance of projections (panel F, white asterisk). Loosening and disorganization of the muscle fibers (panel F and H, lm and cm) and uncommon projections of the spines in the outer tegument (panel H, white asterisks) are observed. Control parasites (panel I) show normal and preserved mitochondria (m), always associated with muscle fibers (lm or cm). MC3935-treated parasites (panel J) lack muscle fibers, and they show smaller mitochondria (m) that appear to have less defined cristae, to be enveloped by membranous structures and to be close to myelin fibers (mf). Scale bars: white (5 µm) and yellow (1 µm).

Large vacuoles were seen in the more external region of the tegument of MC3935-treated parasites (tv in panel D). Additionally, these internal vacuoles contained what seemed to be cellular debris (panel D, #), which could be an indication of tissue degradation and cell death. Panel F shows significant thickening and higher electron density of the outer tegument, associated with the appearance of projections (white asterisk). Loosening and disorganization of the muscle fibers were also observed (panel F and H, lm, and cm) as were different projections of the spines in the outer tegument (panel H, white asterisks). Control parasites (panel I) showed normal and preserved mitochondria (m), always associated with muscle fibers while MC3935-treated parasites (panel J) showed smaller mitochondria (m) that appeared to have less well-defined cristae, and enveloped by membranous structures, which could be an indication of leftover muscle fibers, and they were close to myelin fibers (mf).

### Phenotypic defects of adult *S. mansoni* after SmLSD1 inhibition

Analysis of the adult male tegument and its oral sucker by scanning electron microscopy showed significant alterations upon MC3935 treatment (Fig 7, right panels), when compared to the control worms (Fig 7, left panels). A detailed inspection of the SEM images revealed extensive damage in the dorsal tegument of the male worms that were treated with the LSD1 inhibitor, with the presence of a large number of blisters (Fig 7B and D, yellow arrowheads), as well as fissures and holes in the tubercles (Fig 7B and D, yellow arrows). Blisters and fissures were also seen in the male oral sucker (Fig 7F, yellow arrows, and arrowheads). Confocal laser scanning microscopy (CLSM) showed important alterations of the sexual organs of MC3935-treated male or female parasites (Fig 7H and J). It is worth noting the deleterious effect of the LSD1 inhibitor in the involution of the ovary, leading to a reduced number of mature or immature oocytes (Fig 7, compare panels G and H). The inhibitor also generated severe disorganization of the testicular lobes, culminating in a significantly reduced number of spermatocytes (Fig 7, compare panels I and J). Since the treatment of MC3935 significantly affected the sexual organs of both male and female worms, one should expect that egg production would be severely compromised. Supplementary videos S3 and S4 display the described phenotypic abnormalities, which explain the egg laying impairment in MC3935-treated worms. Indeed, inspection of egg laying by the worms cultivated in the presence of MC3935 revealed an almost complete lack of eggs (Supplementary Fig S4A and B and Supplementary videos S3 and S4; in the videos, note the presence of eggs in DMSO-treated worms and a complete lack of eggs in MC3935-treated parasites) a few hours after the addition of the inhibitor. It is also worth noting the involution of the vitellaria in MC3935-treated females (Supplementary video S4).

**Figure 7.**
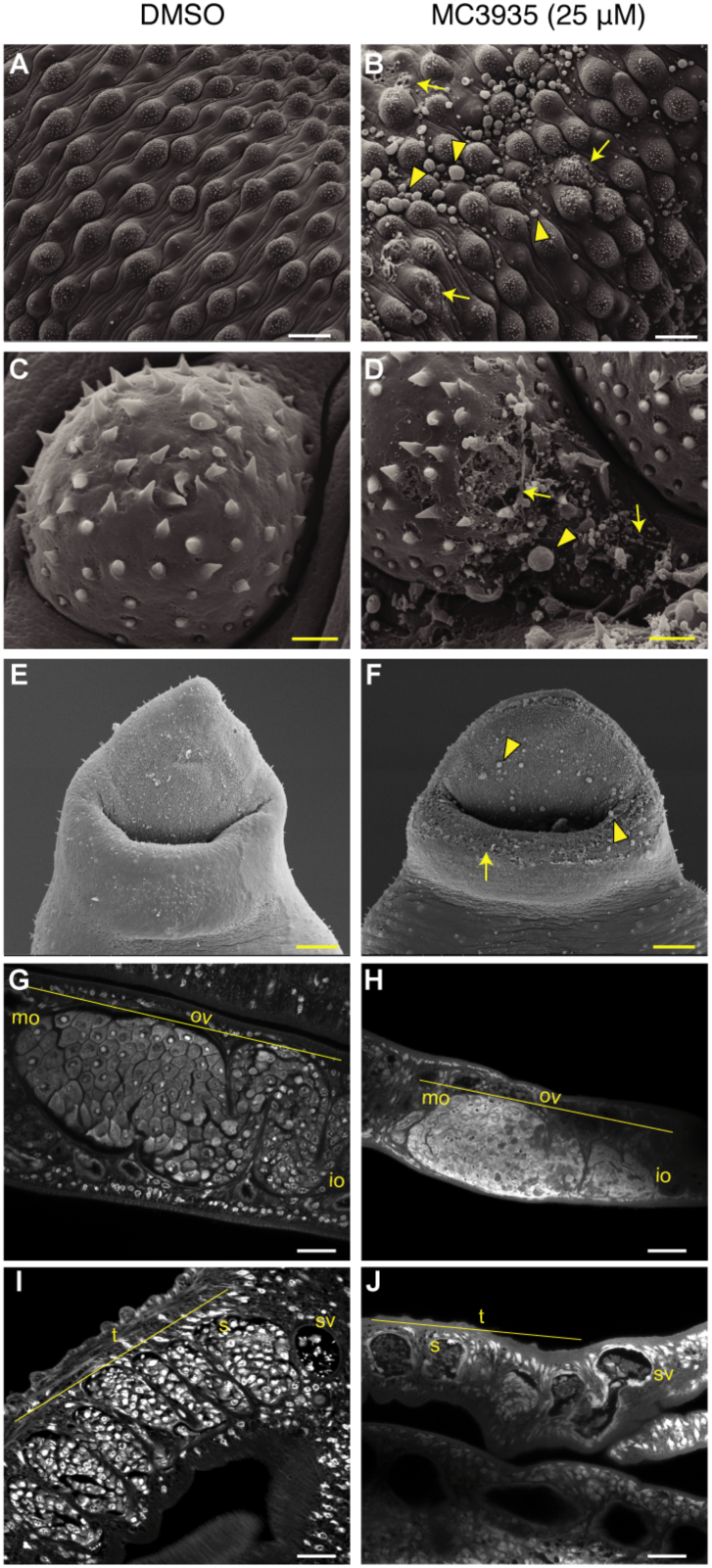
Inhibition of SmLSD1 leads to tegumental damage and reproductive organ involution in adult schistosomes. Ten adult worm pairs were cultivated in the presence of DMSO (left column) or 25 µM MC3935 (right column) for 72 h. Scanning electron microscopy (SEM) images from the dorsal region of male worms show damage to the tegument (B), tuberculous (D) and oral sucker (F) compared to controls (A, C and E). Yellow arrows point to fissures and arrowheads point to blisters. Scale bar: 2 µm (yellow). Confocal laser scanning microscopy (CLSM) of ovaries (G and H) and testis (I and J) from control and MC3935-treated, respectively. OV: ovary; mo: mature oocytes; io: imature oocytes; (t) testicular lobes; sv: seminal vesicle; s: spermocytes. Scale bars: 20 µm (white) and 2 µm (yellow).

### Changes in gene expression profile upon SmLSD1 inhibition

We performed RNA sequencing (RNA-Seq) analysis to evaluate the effect of LSD1 inhibition on global gene transcription in *S. mansoni*. Unsupervised hierarchical clustering analysis of RNA-seq data depicted the changes in global gene expression profile in males, females, or schistosomula upon treatment with MC3935 (Fig 8). Interestingly, inhibition of SmLSD1 significantly modulated 3608 transcribed genes in male, female or schistosomula, with 1964 being downregulated, and 1644 being upregulated (Fig 8A). The highest modulation of gene expression was observed in male worms, for either up- or downregulation (Fig 8A, purple in the Venn diagram), followed by female worms (Fig 8A, pink in the Venn diagram), and schistosomula (Fig 8A, red in the Venn diagram). Importantly, when we analyzed commonly regulated genes in male, female, or schistosomula, we found 220 and 50 genes down- or upregulated, respectively (Fig 8A). The complete lists of significantly upregulated genes in each female, male or schistosomula (Tables S1-S3), downregulated genes in each females, males or schistosomula (Tables S4-S6), down- or upregulated genes in common between females and males (Tables S7 and S8, respectively) are presented in the supporting information. It is noteworthy that in schistosomula, a smaller number of consensus of differentially expressed genes (DEGs) was detected when compared with adult worms (Fig. 8A).

**Figure 8.**
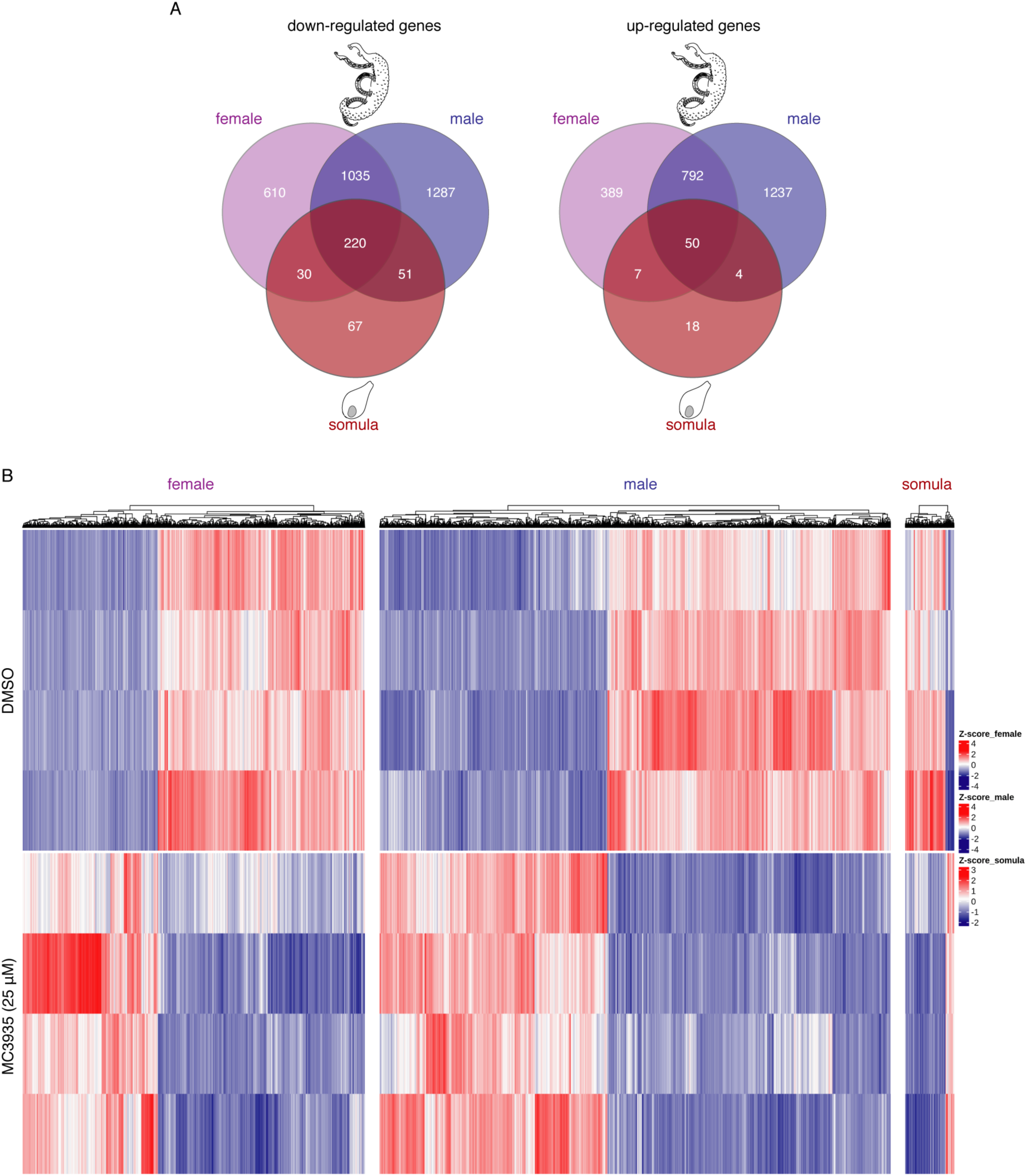
Inhibition of SmLSD1 triggers genome-wide transcriptional deregulation. RNA-seq analysis of female and male worms and schistosomula that were cultivated in the presence of DMSO (control) or MC3935 (25 µM) for 48 hours. (A) Venn diagrams of the number of genes that were detected as differentially expressed among female, male and schistosomula. The numbers at the intersections (darkest red) of the diagrams represent genes commonly affected in the presence of MC3935 among females, males and schistosomula (220 downregulated genes and 50 upregulated genes). (B) The heatmaps show the hierarchical clustering of differentially expressed genes (columns) in four biological replicates (lines) of female and male worms, and schistosomula, either for controls or for treated parasites, as indicated on the left side of the heatmaps. Blue lines, downregulated genes. Red lines, upregulated genes. Gene expression levels are shown as Z-scores, which represent the number of standard deviations above (red) or below (blue) the mean expression value among treated and control samples for each gene; the expression level Z-scores are color-coded as indicated on the scale at the right side of the heatmap.

A clear profile of the differential gene expression between control and MC3935-treated parasites is depicted in the heatmap (Fig. 8B), which confirmed that the treatment led to a significant change in the regulation of genes in females, males and schistosomula, with many genes being either up- (Fig. 8B, red) or downregulated (Fig. 8B, blue).

A closer look at the twenty most differentially expressed genes in females, males or schistosomula revealed that inhibition of SmLSD1 by MC3935 changed the levels of expression of genes belonging to different critical biological processes, including protein and lipid degradation, RNA processing, calcium and sodium homeostasis, antioxidants and transcription factors (Tables S9-S11). Within this list, genes encoding proteases stood out as being downregulated in MC3935-treated schistosomula, males and females as compared to DMSO-treated parasites (Tables S9-S11, shaded in green). Genes encoding protease inhibitors were upregulated in females and males (Tables S9 and S10, shaded in dark green). Importantly, genes of the digestive system of *S. mansoni* were found down- or upregulated (Tables S9-S11, shaded in green with *).

Genes encoding kinases and phosphatases were upregulated in treated females and males (Tables S9 and S10, shaded pink and yellow, respectively). Overall, the analysis of the twenty most differentially expressed genes in treated males revealed a more heterogeneous gene expression profile (Table S10). Of note, in MC3935-treated schistosomula, a significant number of genes encoding proteins involved in RNA metabolism were upregulated (Table S11, shaded in blue). Downregulated genes encoding proteases stood out in treated schistosomula (Table S11, shaded in green).

### Knockdown of the SmLSD1 gene

We conducted dsRNAi experiments in adult worm pairs and were able to achieve 70% silencing of SmLSD1 transcription at day 7 (Supplementary Fig S6A and B). Western blot analysis of protein extracts from silenced adult worm pairs revealed a 2-fold inhibition of SmLSD1 demethylase activity (Supplementary Fig S6C). By monitoring the behavior of the worms on a daily basis, we clearly observed progressive harm in whole-worm physiology during SmLSD1 silencing, which culminated in significant unpairing of male and female worms, a decrease of nearly 50% in the number of laid eggs, low motility and compromised surface adherence by the oral sucker of male worms (Supplementary Fig S6D - F). Silencing of SmLSD1 promoted damage to the oral sucker of male worms (Supplementary Fig S6G). Of note, the silencing of the parasites with the control dsRNAi showed no decrease in SmLSD1 gene expression or activity and no deleterious effects in the worms (Supplementary Fig S6, dsGFP)

## Discussion

Lysine-specific demethylase 1 (LSD1) is an epigenetic enzyme that oxidatively cleaves methyl groups from mono and dimethyl Lysine 4 of histone H3 (H3K4me1/2) and can contribute to gene silencing (38–41). Since its discovery, LSD1 histone demethylase activity has been investigated as a pharmacologic target for cancer and other diseases. As part of a continuing effort of several different investigators to identify epigenetic modifications as therapeutic targets to control schistosomiasis, a recent paper published during our study also identified *S. mansoni* LSD1 (SmLSD1) as an additional promising drug candidate (17). In the present study, we provide evidence that SmLSD1 plays important biological roles in the physiology of *S. mansoni* and that inhibition of SmLSD1 by a novel, selective, and potent LSD1 synthetic inhibitor MC3935 is detrimental to the survival of juvenile and adult worms.

MC3935 was synthesized based on the scaffold of the nonselective and irreversible monoamine oxidase (MAO) inhibitor tranylcypromine (TCP) (29, 42). Tranylcypromine is a mechanism-based suicide inhibitor of MAO and LSD1; it covalently binds to the FAD cofactor embedded within the protein, thus abolishing enzymatic catalysis (43). Importantly, MC3935 showed a 1,000 fold higher inhibitory activity than TCP, *in vitro*. In addition, our *in silico* molecular docking indicated that MC3935 could indeed be an effective SmLSD1 inhibitor and that the inhibition also relied on the amino-oxidase-like catalytic domain. Toxicity assays on *S. mansoni* parasites using each of our 11 synthetic putative LSD1 inhibitors, as well as TCP, revealed MC3935 as the most powerful, with TCP showing the lowest toxicity toward the worms. This was a surprising result if we consider that both MC3935 and TCP (44) should adopt a similar orientation in their binding to the catalytic site of SmLSD1, based on our *in silico* molecular docking. However, one could always assume that the lack of toxicity of TCP was due to its lower permeability to the tegument of *S. mansoni* or that it is more metabolically labile. Indeed, MC3935 fills better the catalytic tuibe of the enzyme than TCP, due to its addiotional ethynylbenzamide portion, which allows additional interaction into the tube. Importantly, our western blot analyses indicated that MC3935 specifically inhibited H3K4 mono- and dimethylation, which are known LSD1 targets, but not H3K4 trimethylation, a mark that is targeted by the histone demethylases of the Jumonji family (doi: 10.1101/gad.1652908).

Genetic studies in numerous models have suggested that LSD1 plays a significant role in developmental processes (45). LSD1 has also been reported to have a role in the DNA damage response (46), repression of mitochondrial metabolism, lipid oxidation energy expenditure programs (47, 48), and smooth muscle regeneration (49). Additionally, germline murine knockouts exhibit embryonic lethality before E7.5: the egg cylinder fails to elongate and gastrulate, resulting in development arrest (50). In *C. elegans*, the homolog spr-5 regulates Notch signaling (51, 52), and maintains transgenerational epigenetic memory and fertility (53). The yeast spLsd1/2 (54) and *Drosophila* Su(var)3-3 (55) homologs regulate gene silencing, which is required to guarantee normal oogenesis (56) and spermatogenesis (57) in *Drosophila*.

Taking into account the phenotypic effects observed in schistosomula or adult worms under the treatment with the LSD1 inhibitor MC3935, it can be assumed that SmLSD1 also plays important roles in the development and homeostasis of the tegument, muscle and sexual organs of *S. mansoni*. Although inhibition of SmLSD1 can have more pronounced effects on many other key biological processes or structures in *S. mansoni*, the tegument disruption would alone represent a desirable LSD1 target. The tegument of *S. mansoni* plays a crucial role in its protection against the host immune system (58); it is capable of absorbing nutrients and molecules, as well as excreting metabolites and synthesizing proteins (59, 60). Inhibition of SmLSD1 activity, either by irreversible MC3935 binding (Fig 3A and B) or by partial knockdown of the SmLSD1 gene (Supplementary Fig S6A and B), generated pronounced damage in schistosomula or adult worms (Fig 5, Fig 7, and Supplementary Fig S6), which likely made a major contribution to the observed mortality of the parasites within a short period of time (Fig 2, Fig 4, Supplementary Fig S3, Supplementary Fig S6, and Supplementary videos).

The musculatory activity is essential in several aspects of *S. mansoni* biology and physiology, including infection, pairing, feeding, regurgitation, reproduction and egg laying (61–63). Our transmission electron microscopy revealed a complete lack of muscle layers in schistosomula treated with the LSD1 inhibitor (Fig 6), which was accompanied by phenotypic defects observed in treated-worms, such as lack of motility, sucker adherence, egg laying and vitellaria contraction (Fig 2, Supplementary Fig S6 and Supplementary video S4), that could be related to the compromised muscle structures.

Interestingly, the role of LSD1 in oogenesis and spermatogenesis seems to be conserved among different organisms, including human, *C. elegans*, *Drosophila,* and *S. mansoni* (this paper). Our confocal microscopy revealed significant alterations in the sexual organs of treated worm pairs, such as a reduced number of spermatocytes and oocytes (Fig 7H and J). These data are in agreement with the incapacity of the worms to produce eggs, even when remaining paired during the treament (see Supplementary Fig S4).

Several diseases have been associated with aberrant histone methylation/demethylation patterns. Thus, considering that juvenile or adult *S. mansoni* are highly transcriptionally active and that SmLSD1 is expected to control the expression of a variety of different genes to maintain the homeostasis of the parasites, we compared their global transcriptional profiles after incubation with a sublethal dose of MC3935. Our RNA-seq analysis revealed that inhibition of SmLSD1 led to the differential expression of genes involved in several different biological processes. These data suggest that SmLSD1 is recruited to target gene promoters by different transcription factors. Among the genes that were downregulated upon SmLSD1 inhibition in schistosomula, females and males, several proteases stood out (Tables S9-S11, shaded in green), mainly from the cathepsin family, including two hemoglobinases in female worms. This is an important finding considering that proteases are key components of the pathogenicity of the parasite; they facilitate tissue penetration and determine the nutritional sources of the parasite within intermediate and human hosts (64). Importantly, two protease inhibitors were upregulated in females and males (Tables S9 and S10, shaded in dark green). That SmLSD1 controls the expression of genes of the digestive system of female or male worms was supported by the fact that a number of genes encoding enzymes or proteins of the digestive tract of the parasite (65) had modified expression in treated parasites (Tables S9-S11, shaded in green and with *). In this regard, it is noteworthy that blood digestion seemed to be severely compromised in females (see Supplementary Video S4).

Interestingly, we identified the SMDR2 gene as significantly downregulated in female worms treated with the LSD1 inhibitor (Table S9, shaded in green and with #). SMDR2 is the schistosome homolog of P-glycoprotein (PgP) (66), an ATP-dependent efflux pump, and its downregulation upon treatment might be involved in the high toxicity of MC3935 observed in females (see Supplementary Video S4).

Importantly, a significant number of proteins involved in RNA metabolism seemed to be specifically upregulated in schistosomula (Table S11, shaded in blue). This finding is of great importance since, although histone methylation has been only recently coupled to RNA processing (67), this is the first report suggesting a role for LSD1 in regulating RNA metabolism.

The epigenetic regulation of chromatin affects many fundamental biological pathways. Therefore, the realization that the deregulation of chromatin is harmful to *S. mansoni* should spark significant efforts to develop selective epigenetic drugs against schistosomiasis. The present study further promotes SmLSD1 as a strong candidate.

## Acknowledgments

We thank Mr. Paulo Cesar dos Santos (Fiocruz, Rio de Janeiro, Brazil) for providing *S. mansoni* cercariae. We are in debt to Dr. Geetha Venkatesh for critical comments and valuable discussions on the bioinformatics analysis.

## Support information Captions

**Supplementary Figure 1. Synthesis and IC_50_ of MC3935.**

**Supplementary Figure 2. *Schistosoma mansoni* LSD1 protein alignment.**

**Supplementary Figure 3. Screening of synthetic small LSD1 inhibitors in *Schistosoma mansoni*.**

**Supplementary Figure 4. LSD1 inhibition affects egg production.**

**Supplementary Figure 5. SmLSD1 inhibition by MC3935 has no effect on H3K4 trimethylation in schistosomes.**

**Supplementary Figure 6. SmLSD1 knockdown partially recapitulates MC3935 phenotypes in adult worms.**

## References

1. Colley DG, Bustinduy AL, Secor WE, King CH. Human schistosomiasis. Lancet [Internet]. 2014 Jun [cited 2019 Sep 5];383(9936):2253–64. Available from: https://linkinghub.elsevier.com/retrieve/pii/S0140673613619492

2. Lewis FA, Tucker MS. Schistosomiasis. In Springer, New York, NY; 2014 [cited 2019 Jun 17]. p. 47–75. Available from: http://link.springer.com/10.1007/978-1-4939-0915-5_3

3. Cioli D, Basso A, Valle C, Pica-Mattoccia L. Decades down the line: the viability of praziquantel for future schistosomiasis treatment. Expert Rev Anti Infect Ther [Internet]. 2012 Aug 10 [cited 2019 Sep 5];10(8):835–7. Available from: http://www.tandfonline.com/doi/full/10.1586/eri.12.70

4. Cioli D, Pica-Mattoccia L, Basso A, Guidi A. Schistosomiasis control: praziquantel forever? Mol Biochem Parasitol [Internet]. 2014 Jun [cited 2019 Sep 5];195(1):23–9. Available from: https://linkinghub.elsevier.com/retrieve/pii/S0166685114000759

5. Pica-Mattoccia L, Doenhoff MJ, Valle C, Basso A, Troiani A-R, Liberti P, et al. Genetic analysis of decreased praziquantel sensitivity in a laboratory strain of Schistosoma mansoni. Acta Trop [Internet]. 2009 Jul [cited 2019 Sep 5];111(1):82–5. Available from: https://linkinghub.elsevier.com/retrieve/pii/S0001706X09000308

6. Liang Y-S, Wang W, Dai J-R, Li H-J, Tao Y-H, Zhang J-F, et al. Susceptibility to praziquantel of male and female cercariae of praziquantel-resistant and susceptible isolates of *Schistosoma mansoni*. J Helminthol [Internet]. 2010 Jun 18 [cited 2019 Sep 5];84(2):202–7. Available from: https://www.cambridge.org/core/product/identifier/S0022149X0999054X/type/journal_article

7. Yang GJ, Lei PM, Wong SY, Ma DL, Leung CH. Pharmacological inhibition of LSD1 for cancer treatment. Vol. 23, Molecules. MDPI AG; 2018.

8. Carneiro VC, de Abreu da Silva IC, Torres EJL, Caby S, Lancelot J, Vanderstraete M, et al. Epigenetic Changes Modulate Schistosome Egg Formation and Are a Novel Target for Reducing Transmission of Schistosomiasis. Knight M, editor. PLoS Pathog [Internet]. 2014 May 8 [cited 2019 Sep 5];10(5):e1004116. Available from: https://dx.plos.org/10.1371/journal.ppat.1004116

9. Marek M, Kannan S, Hauser A-T, Moraes Mourão M, Caby S, Cura V, et al. Structural Basis for the Inhibition of Histone Deacetylase 8 (HDAC8), a Key Epigenetic Player in the Blood Fluke Schistosoma mansoni. Geary TG, editor. PLoS Pathog [Internet]. 2013 Sep 26 [cited 2019 Sep 5];9(9):e1003645. Available from: https://dx.plos.org/10.1371/journal.ppat.1003645

10. Simoben C, Robaa D, Chakrabarti A, Schmidtkunz K, Marek M, Lancelot J, et al. A Novel Class of Schistosoma mansoni Histone Deacetylase 8 (HDAC8) Inhibitors Identified by Structure-Based Virtual Screening and In Vitro Testing. Molecules [Internet]. 2018 Mar 2 [cited 2019 Sep 5];23(3):566. Available from: http://www.mdpi.com/1420-3049/23/3/566

11. Bayer T, Chakrabarti A, Lancelot J, Shaik TB, Hausmann K, Melesina J, et al. Synthesis, Crystallization Studies, and in vitro Characterization of Cinnamic Acid Derivatives as Sm HDAC8 Inhibitors for the Treatment of Schistosomiasis. ChemMedChem [Internet]. 2018 Aug 10 [cited 2019 Sep 5];13(15):1517–29. Available from: http://doi.wiley.com/10.1002/cmdc.201800238

12. Monaldi D, Rotili D, Lancelot J, Marek M, Wössner N, Lucidi A, et al. Structure– Reactivity Relationships on Substrates and Inhibitors of the Lysine Deacylase Sirtuin 2 from Schistosoma mansoni (Sm Sirt2). J Med Chem. 2019;

13. Chucholl C, Morawetz K, Groß H. The clones are coming – strong increase in Marmorkrebs [Procambarus fallax (Hagen, 1870) f. virginalis] records from Europe. Aquat Invasions [Internet]. 2012 [cited 2019 Jun 12];7:511–9. Available from: http://dx.doi.org/10.3391/ai.2012.7.4.008

14. Pereira ASA, Amaral MS, Vasconcelos EJR, Pires DS, Asif H, daSilva LF, et al. Inhibition of histone methyltransferase EZH2 in Schistosoma mansoni in vitro by GSK343 reduces egg laying and decreases the expression of genes implicated in DNA replication and noncoding RNA metabolism. Greenberg RM, editor. PLoS Negl Trop Dis [Internet]. 2018 Oct 26 [cited 2019 Sep 5];12(10):e0006873. Available from: http://dx.plos.org/10.1371/journal.pntd.0006873

15. Roquis D, Taudt A, Geyer KK, Padalino G, Hoffmann KF, Holroyd N, et al. Histone methylation changes are required for life cycle progression in the human parasite Schistosoma mansoni. Hsieh MH, editor. PLOS Pathog [Internet]. 2018 May 21 [cited 2019 Sep 5];14(5):e1007066. Available from: https://dx.plos.org/10.1371/journal.ppat.1007066

16. Cosseau C, Wolkenhauer O, Padalino G, Geyer KK, Hoffmann KF, Grunau C. (Epi)genetic Inheritance in Schistosoma mansoni: A Systems Approach. Trends Parasitol [Internet]. 2017 Apr [cited 2019 Sep 5];33(4):285–94. Available from: https://linkinghub.elsevier.com/retrieve/pii/S1471492216302240

17. Padalino G, Ferla S, Brancale A, Chalmers IW, Hoffmann KF. Combining bioinformatics, cheminformatics, functional genomics and whole organism approaches for identifying epigenetic drug targets in Schistosoma mansoni. Int J Parasitol Drugs Drug Resist [Internet]. 2018 Dec [cited 2019 Sep 5];8(3):559–70. Available from: https://linkinghub.elsevier.com/retrieve/pii/S2211320718301337

18. Maiques-Diaz A, Somervaille TC. LSD1: biologic roles and therapeutic targeting. Epigenomics [Internet]. 2016 Aug [cited 2019 Sep 5];8(8):1103–16. Available from: https://www.futuremedicine.com/doi/10.2217/epi-2016-0009

19. Aravind L, Iyer LM. The SWIRM domain: a conserved module found in chromosomal proteins points to novel chromatin-modifying activities. Genome Biol [Internet]. 2002 Jul 24 [cited 2019 Oct 15];3(8):RESEARCH0039. Available from: http://www.ncbi.nlm.nih.gov/pubmed/12186646

20. Stavropoulos P, Blobel G, Hoelz A. Crystal structure and mechanism of human lysine-specific demethylase-1. Nat Struct Mol Biol [Internet]. 2006 Jul 25 [cited 2019 Sep 5];13(7):626–32. Available from: http://www.nature.com/articles/nsmb1113

21. Shi Y, Lan F, Matson C, Mulligan P, Whetstine JR, Cole PA, et al. Histone Demethylation Mediated by the Nuclear Amine Oxidase Homolog LSD1. Cell [Internet]. 2004 Dec [cited 2019 Sep 5];119(7):941–53. Available from: https://linkinghub.elsevier.com/retrieve/pii/S0092867404011997

22. Lee A, Borrello MT, Ganesan A. LSD (Lysine-Specific Demethylase): A Decade-Long Trip from Discovery to Clinical Trials. In 2019. p. 221–61.

23. Pierce R. Targeting Schistosome Histone Modifying Enzymes for Drug Development. Curr Pharm Des. 2012;

24. Ciccarelli FD, Doerks T, von Mering C, Creevey CJ, Snel B, Bork P. Toward Automatic Reconstruction of a Highly Resolved Tree of Life. Science (80-) [Internet]. 2006 Mar 3 [cited 2019 Oct 15];311(5765):1283–7. Available from: http://www.ncbi.nlm.nih.gov/pubmed/16513982

25. Rotili D, Tomassi S, Conte M, Benedetti R, Tortorici M, Ciossani G, et al. Pan-Histone Demethylase Inhibitors Simultaneously Targeting Jumonji C and Lysine-Specific Demethylases Display High Anticancer Activities. J Med Chem [Internet]. 2014 Jan 9 [cited 2019 Sep 5];57(1):42–55. Available from: https://pubs.acs.org/doi/10.1021/jm4012802

26. Ourailidou ME, Lenoci A, Zwergel C, Rotili D, Mai A, Dekker FJ. Towards the development of activity-based probes for detection of lysine-specific demethylase-1 activity. Bioorg Med Chem [Internet]. 2017 Feb [cited 2019 Sep 5];25(3):847–56. Available from: https://linkinghub.elsevier.com/retrieve/pii/S0968089616312640

27. Bateman A, Martin MJ, O’Donovan C, Magrane M, Alpi E, Antunes R, et al. UniProt: the universal protein knowledgebase. Nucleic Acids Res [Internet]. 2017 Jan 4 [cited 2019 Oct 15];45(D1):D158–69. Available from: https://academic.oup.com/nar/article-lookup/doi/10.1093/nar/gkw1099

28. Webb B, Sali A. Comparative Protein Structure Modeling Using MODELLER. In: Current Protocols in Bioinformatics [Internet]. Hoboken, NJ, USA: John Wiley & Sons, Inc.; 2016 [cited 2019 Oct 15]. p. 5.6.1-5.6.37. Available from: http://doi.wiley.com/10.1002/cpbi.3

29. Binda C, Valente S, Romanenghi M, Pilotto S, Cirilli R, Karytinos A, et al. Biochemical, Structural, and Biological Evaluation of Tranylcypromine Derivatives as Inhibitors of Histone Demethylases LSD1 and LSD2. J Am Chem Soc [Internet]. 2010 May 19 [cited 2019 Oct 15];132(19):6827–33. Available from: http://www.ncbi.nlm.nih.gov/pubmed/20415477

30. Smithers SR, Terry RJ. The infection of laboratory hosts with cercariae of *Schistosoma mansoni* and the recovery of the adult worms. Parasitology [Internet]. 1965 Nov 10 [cited 2019 Jun 17];55(4):695–700. Available from: https://www.cambridge.org/core/product/identifier/S0031182000086248/type/journal_article

31. Lancelot J, Caby S, Dubois-Abdesselem F, Vanderstraete M, Trolet J, Oliveira G, et al. Schistosoma mansoni Sirtuins: Characterization and Potential as Chemotherapeutic Targets. Williams DL, editor. PLoS Negl Trop Dis [Internet]. 2013 Sep 12 [cited 2019 Jun 17];7(9):e2428. Available from: https://dx.plos.org/10.1371/journal.pntd.0002428

32. Panic G, Flores D, Ingram-Sieber K, Keiser J. Fluorescence/luminescence-based markers for the assessment of Schistosoma mansoni schistosomula drug assays. Parasites and Vectors. 2015 Dec 8;8(1).

33. Dubois F, Caby S, Oger F, Cosseau C, Capron M, Grunau C, et al. Histone deacetylase inhibitors induce apoptosis, histone hyperacetylation and up-regulation of gene transcription in Schistosoma mansoni. Mol Biochem Parasitol [Internet]. 2009 Nov [cited 2019 Sep 5];168(1):7–15. Available from: https://linkinghub.elsevier.com/retrieve/pii/S0166685109001601

34. Carneiro VC, de Abreu da Silva IC, Torres EJL, Caby S, Lancelot J, Vanderstraete M, et al. Epigenetic Changes Modulate Schistosome Egg Formation and Are a Novel Target for Reducing Transmission of Schistosomiasis. PLoS Pathog. 2014;10(5).

35. Storey DM, Ogbogu VC. Observations on third-stage larvae and adults of Litomosoides carinii (Nematoda: Filarioidea) by scanning and transmission electron microscopy. Ann Trop Med Parasitol. 1991;85(1):111–21.

36. Pereira ASA, Amaral MS, Vasconcelos EJR, Pires DS, Asif H, daSilva LF, et al. Inhibition of histone methyltransferase EZH2 in Schistosoma mansoni in vitro by GSK343 reduces egg laying and decreases the expression of genes implicated in DNA replication and noncoding RNA metabolism. Greenberg RM, editor. PLoS Negl Trop Dis [Internet]. 2018 Oct 26 [cited 2019 Jun 17];12(10):e0006873. Available from: http://dx.plos.org/10.1371/journal.pntd.0006873

37. Bolger AM, Lohse M, Usadel B. Trimmomatic: a flexible trimmer for Illumina sequence data. Bioinformatics [Internet]. 2014 Aug 1 [cited 2019 Sep 5];30(15):2114–20. Available from: https://academic.oup.com/bioinformatics/article-lookup/doi/10.1093/bioinformatics/btu170

38. Burg JM, Link JE, Morgan BS, Heller FJ, Hargrove AE, McCafferty DG. KDM1 class flavin-dependent protein lysine demethylases. Biopolymers [Internet]. 2015 Jul [cited 2019 Oct 15];104(4):213–46. Available from: http://www.ncbi.nlm.nih.gov/pubmed/25787087

39. Bennesch MA, Segala G, Wider D, Picard D. LSD1 engages a corepressor complex for the activation of the estrogen receptor α by estrogen and cAMP. Nucleic Acids Res [Internet]. 2016 Oct 14 [cited 2019 Oct 15];44(18):8655–70. Available from: http://www.ncbi.nlm.nih.gov/pubmed/27325688

40. Pilotto S, Speranzini V, Tortorici M, Durand D, Fish A, Valente S, et al. Interplay among nucleosomal DNA, histone tails, and corepressor CoREST underlies LSD1-mediated H3 demethylation. Proc Natl Acad Sci [Internet]. 2015 Mar 3 [cited 2019 Oct 15];112(9):2752–7. Available from: http://www.ncbi.nlm.nih.gov/pubmed/25730864

41. Dimitrova E, Turberfield AH, Klose RJ. Histone demethylases in chromatin biology and beyond. EMBO Rep [Internet]. 2015 Dec 12 [cited 2019 Oct 15];16(12):1620–39. Available from: http://www.ncbi.nlm.nih.gov/pubmed/26564907

42. Frieling H, Bleich S. Tranylcypromine. Eur Arch Psychiatry Clin Neurosci [Internet]. 2006 Aug [cited 2019 Oct 15];256(5):268–73. Available from: http://www.ncbi.nlm.nih.gov/pubmed/16927039

43. Mimasu S, Sengoku T, Fukuzawa S, Umehara T, Yokoyama S. Crystal structure of histone demethylase LSD1 and tranylcypromine at 2.25Å. Biochem Biophys Res Commun [Internet]. 2008 Feb 1 [cited 2019 Oct 15];366(1):15–22. Available from: http://www.ncbi.nlm.nih.gov/pubmed/18039463

44. Lee MG, Wynder C, Schmidt DM, McCafferty DG, Shiekhattar R. Histone H3 Lysine 4 Demethylation Is a Target of Nonselective Antidepressive Medications. Chem Biol [Internet]. 2006 Jun [cited 2019 Oct 15];13(6):563–7. Available from: http://www.ncbi.nlm.nih.gov/pubmed/16793513

45. Shi Y. Histone lysine demethylases: emerging roles in development, physiology and disease. Nat Rev Genet [Internet]. 2007 Nov [cited 2019 Oct 15];8(11):829–33. Available from: http://www.ncbi.nlm.nih.gov/pubmed/17909537

46. Mosammaparast N, Kim H, Laurent B, Zhao Y, Lim HJ, Majid MC, et al. The histone demethylase LSD1/KDM1A promotes the DNA damage response. J Cell Biol [Internet]. 2013 Nov 11 [cited 2019 Oct 15];203(3):457–70. Available from: http://www.ncbi.nlm.nih.gov/pubmed/24217620

47. Hino S, Sakamoto A, Nagaoka K, Anan K, Wang Y, Mimasu S, et al. FAD-dependent lysine-specific demethylase-1 regulates cellular energy expenditure. Nat Commun [Internet]. 2012 Jan 27 [cited 2019 Oct 15];3(1):758. Available from: http://www.nature.com/articles/ncomms1755

48. Sakamoto A, Hino S, Nagaoka K, Anan K, Takase R, Matsumori H, et al. Lysine Demethylase LSD1 Coordinates Glycolytic and Mitochondrial Metabolism in Hepatocellular Carcinoma Cells. Cancer Res [Internet]. 2015 Apr 1 [cited 2019 Oct 15];75(7):1445–56. Available from: http://www.ncbi.nlm.nih.gov/pubmed/25649769

49. Tosic M, Allen A, Willmann D, Lepper C, Kim J, Duteil D, et al. Lsd1 regulates skeletal muscle regeneration and directs the fate of satellite cells. Nat Commun [Internet]. 2018 Dec 25 [cited 2019 Oct 15];9(1):366. Available from: http://www.nature.com/articles/s41467-017-02740-5

50. Wang J, Scully K, Zhu X, Cai L, Zhang J, Prefontaine GG, et al. Opposing LSD1 complexes function in developmental gene activation and repression programmes. Nature [Internet]. 2007 Apr 28 [cited 2019 Oct 15];446(7138):882–7. Available from: http://www.ncbi.nlm.nih.gov/pubmed/17392792

51. Jarriault S, Greenwald I. Suppressors of the egg-laying defective phenotype of sel-12 presenilin mutants implicate the CoREST corepressor complex in LIN-12/Notch signaling in C. elegans. Genes Dev [Internet]. 2002 Oct 15 [cited 2019 Oct 15];16(20):2713–28. Available from: http://www.ncbi.nlm.nih.gov/pubmed/12381669

52. Eimer S, Lakowski B, Donhauser R, Baumeister R. Loss of spr-5 bypasses the requirement for the C.elegans presenilin sel-12 by derepressing hop-1. EMBO J [Internet]. 2002 Nov 1 [cited 2019 Oct 15];21(21):5787–96. Available from: http://www.ncbi.nlm.nih.gov/pubmed/12411496

53. Katz DJ, Edwards TM, Reinke V, Kelly WG. A C. elegans LSD1 Demethylase Contributes to Germline Immortality by Reprogramming Epigenetic Memory. Cell [Internet]. 2009 Apr 17 [cited 2019 Oct 15];137(2):308–20. Available from: http://www.ncbi.nlm.nih.gov/pubmed/19379696

54. Lan F, Zaratiegui M, Villén J, Vaughn MW, Verdel A, Huarte M, et al. S. pombe LSD1 Homologs Regulate Heterochromatin Propagation and Euchromatic Gene Transcription. Mol Cell [Internet]. 2007 Apr 13 [cited 2019 Oct 15];26(1):89–101. Available from: http://www.ncbi.nlm.nih.gov/pubmed/17434129

55. Rudolph T, Yonezawa M, Lein S, Heidrich K, Kubicek S, Schäfer C, et al. Heterochromatin Formation in Drosophila Is Initiated through Active Removal of H3K4 Methylation by the LSD1 Homolog SU(VAR)3-3. Mol Cell [Internet]. 2007 Apr 13 [cited 2019 Oct 15];26(1):103–15. Available from: http://www.ncbi.nlm.nih.gov/pubmed/17434130

56. Yang F, Quan Z, Huang H, He M, Liu X, Cai T, et al. Ovaries absent links dLsd1 to HP1a for local H3K4 demethylation required for heterochromatic gene silencing. Elife [Internet]. 2019 Jan 16 [cited 2019 Oct 15];8. Available from: https://elifesciences.org/articles/40806

57. Zhang J, Bonasio R, Strino F, Kluger Y, Holloway JK, Modzelewski AJ, et al. SFMBT1 functions with LSD1 to regulate expression of canonical histone genes and chromatin-related factors. Genes Dev [Internet]. 2013 Apr 1 [cited 2019 Oct 15];27(7):749–66. Available from: http://www.ncbi.nlm.nih.gov/pubmed/23592795

58. Shaw MK, Erasmus DA. Schistosoma mansoni: praziquantel-induced changes to the female reproductive system. Exp Parasitol [Internet]. 1988 Feb [cited 2019 Oct 15];65(1):31–42. Available from: http://www.ncbi.nlm.nih.gov/pubmed/3338547

59. Bertão HG, da Silva RAR, Padilha RJR, de Azevedo Albuquerque MCP, Rádis-Baptista G. Ultrastructural analysis of miltefosine-induced surface membrane damage in adult Schistosoma mansoni BH strain worms. Parasitol Res [Internet]. 2012 Jun 4 [cited 2019 Oct 15];110(6):2465–73. Available from: http://www.ncbi.nlm.nih.gov/pubmed/22215191

60. Xavier AML, Tavares D, Guimarães EV, Sarro-Silva M de F, Silva AC, de Moraes Neto AHA. Ultrastructural alterations in adult Schistosoma mansoni, harbored in non-antihelminthic treated and low-inflammatory mice by transmission electron microscopy (TEM). Acta Trop [Internet]. 2014 Feb [cited 2019 Oct 15];130:51–7. Available from: http://www.ncbi.nlm.nih.gov/pubmed/24161877

61. Ressurreição M, De Saram P, Kirk RS, Rollinson D, Emery AM, Page NM, et al. Protein Kinase C and Extracellular Signal-Regulated Kinase Regulate Movement, Attachment, Pairing and Egg Release in Schistosoma mansoni. Gasser RB, editor. PLoS Negl Trop Dis [Internet]. 2014 Jun 12 [cited 2019 Oct 15];8(6):e2924. Available from: http://www.ncbi.nlm.nih.gov/pubmed/24921927

62. Blair KL, Bennett JL, Pax RA. Schistosoma mansoni: Evidence for protein kinase-C-like modulation of muscle activity. Exp Parasitol [Internet]. 1988 Aug [cited 2019 Oct 15];66(2):243–52. Available from: http://www.ncbi.nlm.nih.gov/pubmed/3165068

63. Mair GR, Maule AG, Day TA, Halton DW. A confocal microscopical study of the musculature of adult Schistosoma mansoni. Parasitology [Internet]. 2000 Aug [cited 2019 Oct 15];121 (Pt 2):163–70. Available from: http://www.ncbi.nlm.nih.gov/pubmed/11085236

64. McKerrow JH. Parasite proteases. Exp Parasitol [Internet]. 1989 Jan [cited 2019 Oct 15];68(1):111–5. Available from: http://www.ncbi.nlm.nih.gov/pubmed/2645160

65. Nawaratna SSK, Gobert GN, Willis C, Chuah C, McManus DP, Jones MK. Transcriptional profiling of the oesophageal gland region of male worms of Schistosoma mansoni. Mol Biochem Parasitol [Internet]. 2014 Sep [cited 2019 Oct 15];196(2):82–9. Available from: http://www.ncbi.nlm.nih.gov/pubmed/25149559

66. Bosch IB, Wang ZX, Tao LF, Shoemaker CB. Two Schistosoma mansoni cDNAs encoding ATP-binding cassette (ABC) family proteins. Mol Biochem Parasitol [Internet]. 1994 Jun [cited 2019 Oct 15];65(2):351–6. Available from: http://www.ncbi.nlm.nih.gov/pubmed/7969275

67. Li Z, Jiang D, Fu X, Luo X, Liu R, He Y. Coupling of histone methylation and RNA processing by the nuclear mRNA cap-binding complex. Nat Plants [Internet]. 2016 Mar 29 [cited 2019 Oct 15];2(3):16015. Available from: http://www.nature.com/articles/nplants201615

