## Supplementary figures for "Pharmacological inhibition of lysine-specific demethylase 1 (LSD1) induces global transcriptional deregulation and ultrastructural alterations that impair viability in *Schistosoma mansoni*"

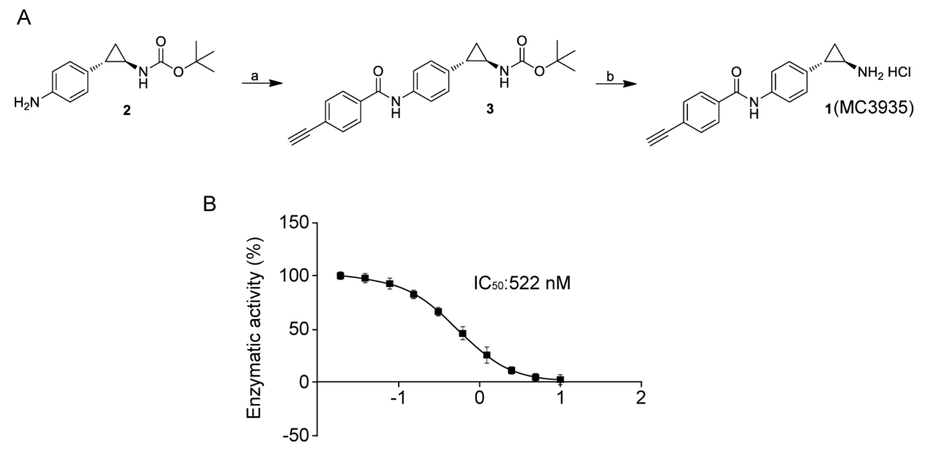


**Supplementary Figure 1. Synthesis and $IC_{50}$ of MC3935.** (A). Compound **1** (MC3935) was synthesized by coupling racemic *tert*-butyl (*trans*-2-(4-(4-ethynylbenzamido)phenyl)cyclopropyl) carbamate **2,** prepared as previously reported,[1] with the commercially available 4-ethynylbenzoic acid followed by acidic deprotection of the Boc protected amine **3.** Reagents for the synthesis of compound **1** (**MC3935**): (a) HOBt, EDCI, TEA, dry DMF, rt; (b) HCl 4N in dioxane, dry THF, 0°C-rt. (B). MC3935 inhibits the catalytic activity of recombinant human LSD1 (hLSD1). The concentration required to inhibit the activity of the purified hLSD1 protein by 50% ($IC_{50}$) is shown in the graph.

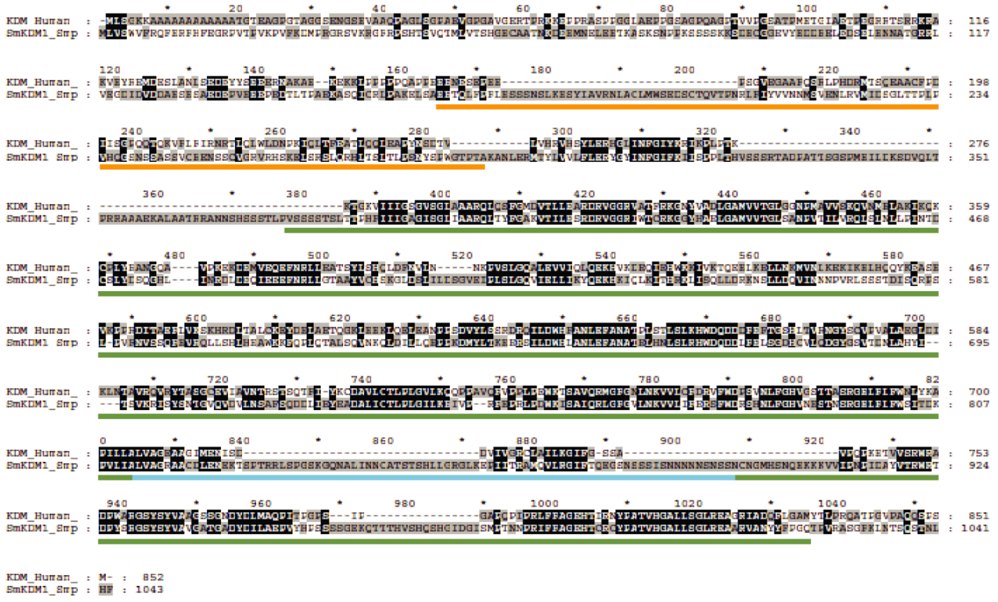

**Supplementary Figure 2.** ***Schistosoma mansoni*** **LSD1 protein alignment.** Sequence alignment using the Clustal Omega tool was performed including the *Homo sapiens* - NP_055828, and *Schistosoma mansoni* – XP_018652619.1. The functional domains of the LSD1 protein family are underlined as follows: the SWIRM domain in orange (165-287 aa), the amino-oxidase-like domain in green (379-824 and 909-1136 aa) and the TOWER domain in blue (825-908 aa). Amino acid positions refer to the SmLSD1 protein. Unique amino acid sequences found within the SmLSD1 polypeptide are shown as dashes.

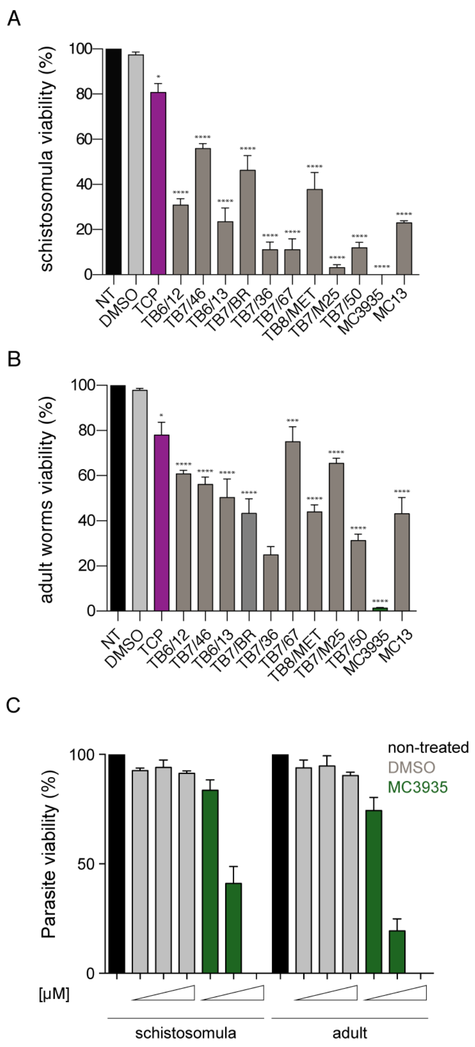


**Supplementary Figure 3. Screening of synthetic small LSD1 inhibitors in *Schistosoma mansoni.*** Twenty thousand schistosomula (A) or ten adult worm pairs (B) were incubated with 25 µM of LSD1 inhibitors or DMSO (nontreated parasites were included as an additional control) and submitted to an ATP cell viability assay. Tranylcypromine (TCP, red bars) is a well-known irreversible LSD1 inhibitor. Twelve different compounds based on the TCP scaffold were tested. (C) Dose-dependent toxicity of MC39350 (at 1, 10 or 25 µM) on schistosomula or adult worm pairs by. Incubation times for schistosomula and adult worms were 72 h and 96 h, respectively. The results of three independent assays are shown; error bars represent the SD.

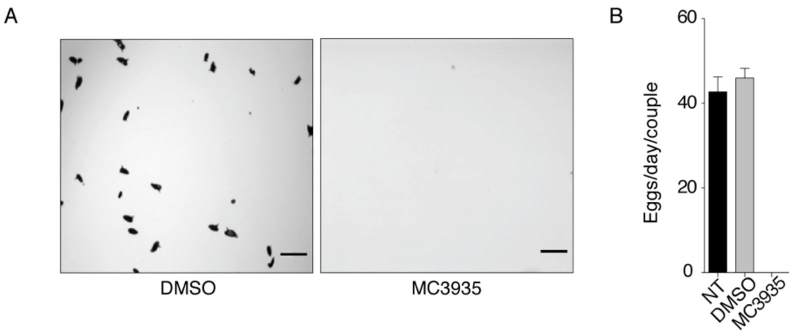


**Supplementary Figure 4. LSD1 inhibition affects egg production.** (A). Adult worm pairs were treated (or not, NT) with 0.25% DMSO or 25 µM MC3935 and cultivated for 96 h. The number of laid eggs was counted daily and a representative image was recorded. Scale bar: 250 µm. (B). Quantification of eggs normalized by the number of adult worm pairs and days of treatment.

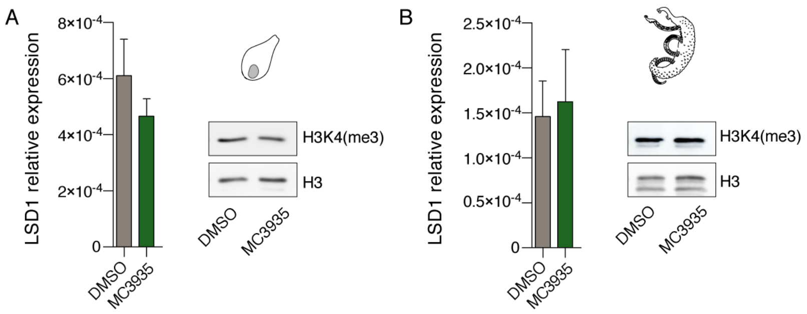


**Supplementary Figure 5. SmLSD1 inhibition by MC3935 has no effect on H3K4 trimethylation in schistosomes.** Quantitative RT-PCR analysis of SmLSD1 mRNA or western blot analyses of SmLSD1 protein from schistosomula (A) or adult worms (B) after 72 h or 96 h incubation time with MC3935, respectively. The bars indicate standard deviations from three independent measurements. Histone H3 was included in western blots as the loading control.

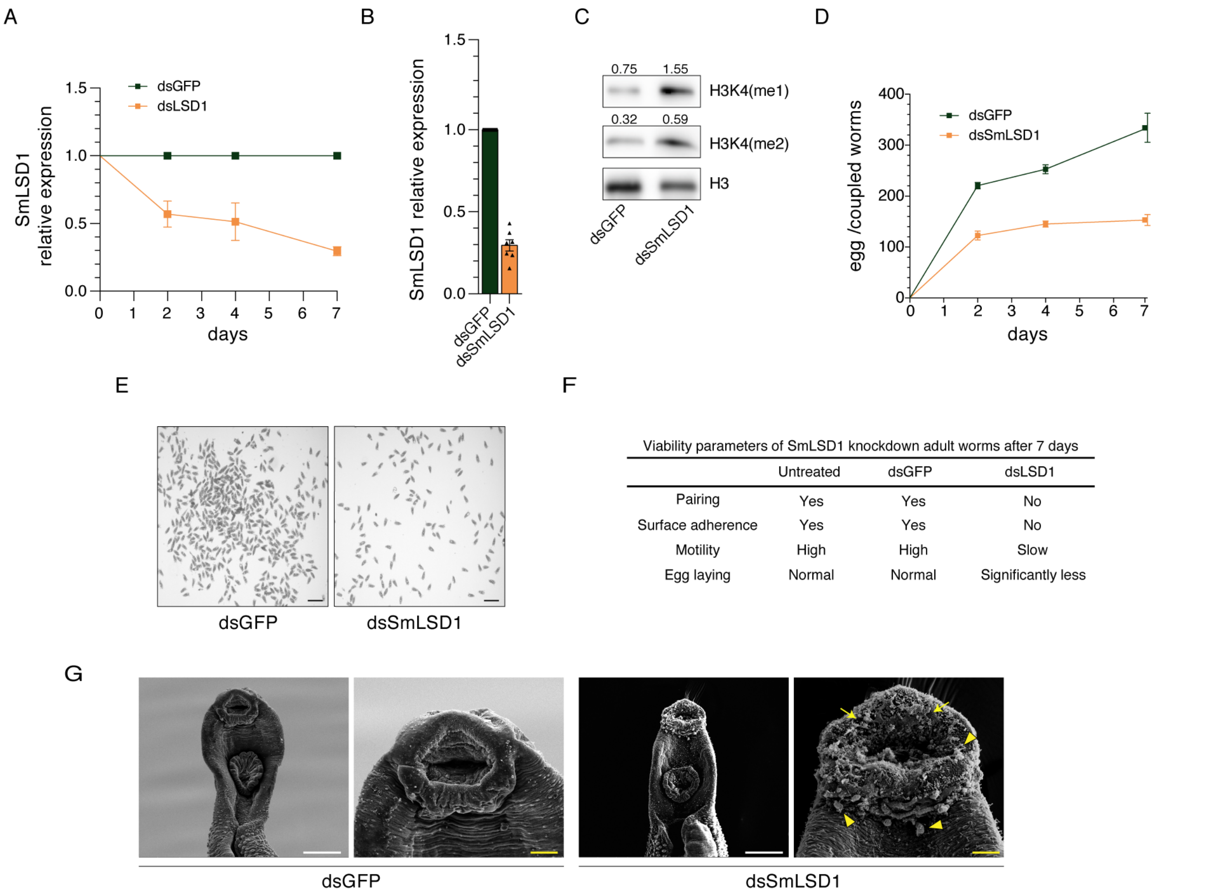


Viability parameters of SmLSD1 knockdown adult worms after 7 days

| | Untreated | dsGFP | dsLSD1 |
|---|---|---|---|
| Pairing | Yes | Yes | No |
| Surface adherence | Yes | Yes | No |
| Motility | High | High | Slow |
| Egg laying | Normal | Normal | Significantly less |

**Supplementary Figure 6. SmLSD1 knockdown partially recapitulates MC3935 phenotypes in adult worms.** (A). Adult worm pairs were soaked with 30 µg of dsDNA and cultivated for up to 7 days. A silencing of 70% was obtained for SmLSD1 mRNA at day 7 (panels A and B). (C) Western blot analysis of total protein extracts from GFP- or SmLSD1-silenced worms at day 7. Band intensity quantifications (obtained with Image J) are shown above each image, and they were normalized by the H3 band. (D and E) Egg production by GFP- or SmLSD1-silenced female worms was monitored daily (scale bar = 200 µm). (F) Several parameters for adult worm viability were monitored daily using a light microscope, until day 7. The viability parameters were reviewed and scored by two independent observers. (G) SmLSD1 RNAi-mediated phenotypic effects observed by scanning electron microscopy. Arrows point to fissures and arrowheads to blisters in the oral sucker of male worms (scale bar = white (5 µm) and yellow (1 µm).

**Supplementary video 1.** Schistosomula treated with DMSO for 48 h.

**Supplementary video 2.** Schistosomula treated with MC3935 for 48 h.

**Supplementary video 3.** Adult worms treated with DMSO for 96 h.

**Supplementary video 4.** Adult worms treated with MC3935 for 96 h.
