## Supplementary tables S9-11 for "Pharmacological inhibition of lysine-specific demethylase 1 (LSD1) induces global transcriptional deregulation and ultrastructural alterations that impair viability in *Schistosoma mansoni*"

### Table S9. MC3935-treated female of *S. mansoni*

| Female downregulated genes | | | | |
|---|---|---|---|---|
| GeneID | Log2FoldChange | | | product_description |
| | Cuffdif | b-Sleuth | EdgeR | |
| Smp_170630 | -3,44 | -2,63 | -3,44 | Periostin 2C putative |
| Smp_085180 | -3,28 | -2,85 | -3,37 | cathepsin B (C01 family)* |
| Smp_155570 | -2,75 | -2,46 | -2,98 | endoglycoceramidase |
| Smp_055780 | -2,84 | -2,61 | -2,96 | smdr2# |
| Smp_174700 | -2,83 | -2,62 | -2,84 | transcription factor HNF 4 |
| Smp_156960 | -2,63 | -2,31 | -2,77 | nardilysin (M16 family) |
| Smp_042720 | -2,77 | -2,02 | -2,67 | beta 13 n galactosyltransferase |
| Smp_187140 | -2,58 | -2,31 | -2,65 | cathepsin L proteinase* |
| Smp_000190 | -2,56 | -2,05 | -2,64 | short chain dehydrogenase:reductase family 16C |
| Smp_147070 | -2,50 | -1,95 | -2,61 | sodium coupled neutral amino acid |
| Smp_169260 | -2,47 | -2,11 | -2,59 | zinc finger protein 362 |
| Smp_166540 | -2,42 | -2,16 | -2,56 | serine:threonine protein kinase Nek11 |
| Smp_166530 | -2,42 | -2,16 | -2,56 | phospholipase A* |
| Smp_085010 | -2,44 | -2,17 | -2,55 | cathepsin B peptidase (C01 family)* |
| Smp_075800 | -2,31 | -2,17 | -2,53 | hemoglobinase (C13 family) |
| Smp_126120 | -2,53 | -2,04 | -2,52 | LAMA protein 2 |
| Smp_036010 | -2,33 | -2,00 | -2,49 | magnesium transporter nipa2 |
| Smp_139160 | -2,53 | -2,27 | -2,48 | SmCL2 peptidase (C01 family) |
| Smp_141030 | -2,25 | -2,28 | -2,46 | epidermal growth factor receptor pathway |
| Smp_016490 | -2,42 | -2,09 | -2,42 | saposin B domain containing protein* |

| Female upregulated genes | | | | |
|---|---|---|---|---|
| GeneID | (Log2FoldChange) | | | product_description |
| | Cuffdif | b-Sleuth | EdgeR | |
| Smp_025390 | 5,76 | 3,84 | 5,77 | calcium dependent protein kinase |
| Smp_128550 | 5,13 | 3,51 | 4,92 | src type protein tyrosine kinase |
| Smp_094930 | 4,76 | 3,02 | 4,66 | early growth response protein 1 |
| Smp_134870 | 4,01 | 2,58 | 3,89 | putative early growth response protein 2 |
| Smp_124060 | 3,15 | 1,97 | 3,02 | venom allergen-like (VAL) 13 protein* |
| Smp_174810 | 3,20 | 1,93 | 2,99 | Extracellular superoxide dismutase (Cu Zn) |
| Smp_159810 | 4,15 | 2,03 | 2,91 | MEG-2 (ESP15) family* |
| Smp_172460 | 2,57 | 1,36 | 2,51 | Krueppel factor 10 like |
| Smp_049300 | 2,44 | 1,73 | 2,50 | major egg antigen 2C putative |
| Smp_186020 | 2,98 | 1,62 | 2,37 | major egg antigen |
| Smp_008660 | 2,31 | 1,48 | 2,31 | gelsolin |
| Smp_135980 | 2,25 | 1,39 | 2,16 | Calcium binding protein |
| Smp_172960 | 2,13 | 1,38 | 2,14 | serine type protease inhibitor |
| Smp_147730 | 2,13 | 1,38 | 2,14 | single kunitz protease inhibitor |
| Smp_034500 | 2,25 | 1,30 | 2,11 | Dual specificity protein phosphatase 10 |
| Smp_053560 | 2,07 | 1,29 | 2,02 | MAP kinase activated protein kinase 2 |
| Smp_123190 | 1,63 | 1,22 | 1,99 | ADP ribosylation factor protein 2 binding |
| Smp_136660 | 2,21 | 1,16 | 1,95 | pro neuregulin 2 |
| Smp_136650 | 2,21 | 1,16 | 1,95 | pro neuregulin 2 membrane bound |
| Smp_028210 | 2,02 | 1,26 | 1,89 | calcyphosin protein like |

### Table S10. MC3935-treated male of *S. mansoni*

| Male downregulated genes | | | | |
|---|---|---|---|---|
| GeneID | (Log2FoldChange) | | | product_description |
| | Cuffdif | b-Sleuth | EdgeR | |
| Smp_180350 | -4,88 | -3,53 | -4,97 | opsin receptor |
| Smp_017620 | -4,90 | -3,39 | -4,83 | membrane primary amine oxidase |
| Smp_017610 | -4,90 | -3,39 | -4,83 | amiloride sensitive amine oxidase |
| Smp_000755 | -4,66 | -3,33 | -4,75 | family M13 non peptidase ue (M13 family) |
| Smp_135230 | -4,62 | -3,30 | -4,62 | Tyrosine DeCarboxylase family member (tdc 1) |
| Smp_160360 | -4,31 | -3,02 | -4,22 | sodium:chloride dependent neurotransmitter |
| Smp_136730 | -4,35 | -3,15 | -4,14 | cathepsin d (lysosomal aspartyl protease)* |
| Smp_149930 | -4,71 | -3,04 | -4,08 | sodium:potassium:calcium exchanger 6 |
| Smp_008610 | -4,01 | -2,91 | -4,06 | deoxyribonuclease ii |
| Smp_104890 | -3,91 | -2,79 | -4,03 | Cys loop ligand gated ion channel subunit |
| Smp_128860 | -4,03 | -2,79 | -3,99 | lysyl oxidase 2 |
| Smp_169570 | -3,95 | -2,76 | -3,94 | glycerol 3 phosphate dehydrogenase |
| Smp_170630 | -3,80 | -2,66 | -3,80 | Periostin 2C putative |
| Smp_165340 | -4,26 | -2,68 | -3,78 | alpha tocopherol transfer protein |
| Smp_143300 | -3,79 | -2,68 | -3,76 | fibrillin 1 |
| Smp_195090 | -3,71 | -2,61 | -3,75 | tegument-allergen-like protein |
| Smp_193350 | -3,69 | -2,55 | -3,63 | cadherin EGF LAG seven pass G type receptor |
| Smp_085180 | -3,77 | -2,68 | -3,59 | cathepsin B (C01 family)* |
| Smp_123780 | -3,62 | -2,58 | -3,56 | glypican 5 |
| Smp_172590 | -3,40 | -2,57 | -3,48 | family S10 unassigned peptidase (S10 family) |

| Male upregulated genes | | | | |
|---|---|---|---|---|
| GeneID | (Log2FoldChange) | | | product_description |
| | Cuffdif | b-Sleuth | EdgeR | |
| Smp_133770 | 5,71 | 3,91 | 5,68 | lengsin |
| Smp_025390 | 4,64 | 3,20 | 4,61 | calcium dependent protein kinase |
| Smp_047680 | 4,21 | 3,09 | 4,41 | ferritin 2C heavy polypeptide 1 |
| Smp_159810 | 4,65 | 2,64 | 3,86 | MEG-2 (ESP15) family* |
| Smp_128550 | 3,17 | 2,28 | 3,37 | src type protein tyrosine kinase |
| Smp_134870 | 3,21 | 2,21 | 3,14 | early growth response protein |
| Smp_172960 | 3,37 | 2,05 | 3,02 | serine type protease inhibitor |
| Smp_147730 | 3,37 | 2,05 | 3,02 | single kunitz protease inhibitor |
| Smp_172460 | 2,98 | 2,05 | 3,01 | Krueppel factor 10 like |
| Smp_048050 | 3,30 | 2,09 | 2,96 | Major egg antigen (p40) |
| Smp_051400 | 2,83 | 1,97 | 2,87 | dynein light chain |
| Smp_210990 | 2,70 | 1,62 | 2,84 | serine:threonine protein phosphatase PP1 beta |
| Smp_049230 | 3,06 | 2,01 | 2,84 | Major egg antigen (p40) |
| Smp_047650 | 2,82 | 1,95 | 2,83 | ferritin 2C heavy polypeptide 1 |
| Smp_049300 | 2,38 | 1,94 | 2,73 | major egg antigen 2C putative |
| Smp_094930 | 2,73 | 1,87 | 2,68 | early growth response protein 1 |
| Smp_129510 | 2,72 | 1,81 | 2,65 | metallophosphoesterase domain containing protein |
| Smp_130370 | 2,51 | 1,27 | 2,47 | elongation of very long chain fatty acids* |
| Smp_077880 | 2,45 | 1,73 | 2,45 | annexin |
| Smp_186020 | 2,65 | 1,56 | 2,43 | major egg antigen |

**Table S11. MC3935-treated schistosomula of *S. mansoni***

| schistosomula downregulated genes | | | | |
|---|---|---|---|---|
| GeneID | (Log2FoldChange) | | | product_description |
| | Cuffdif | b-Sleuth | EdgeR | |
| Smp_169190 | -3,46 | -2,21 | -3,40 | tegument-allergen-like protein |
| Smp_187140 | -2,25 | -1,63 | -2,33 | cathepsin L proteinase* |
| Smp_162770 | -2,18 | -1,58 | -2,30 | lysosome associated membrane glycoprotein |
| Smp_140610 | -1,97 | -1,58 | -2,29 | iron:zinc purple acid phosphatase protein |
| Smp_085010 | -2,15 | -1,56 | -2,22 | cathepsin B peptidase (C01 family)* |
| Smp_105420 | -2,14 | -1,57 | -2,17 | Saposin%2CIPR008139 Saposin |
| Smp_028870 | -2,05 | -1,62 | -2,17 | Zinc finger%2C C2H2 type domain containing protein |
| Smp_067060 | -1,96 | -1,48 | -2,16 | cathepsin B peptidase (C01 family)* |
| Smp_166540 | -2,03 | -1,52 | -2,13 | serine:threonine protein kinase |
| Smp_105450 | -2,20 | -1,51 | -2,12 | saposin containing protein* |
| Smp_020080 | -1,91 | -1,34 | -2,00 | GTP binding protein (I) alpha subunit alpha |
| Smp_032980 | -1,94 | -1,34 | -1,95 | calmodulin protein |
| Smp_139240 | -1,87 | -1,32 | -1,81 | cathepsin S (C01 family) |
| Smp_138270 | -1,85 | -1,20 | -1,79 | diaminopimelate epimerase DafE |
| Smp_142490 | -1,64 | -1,22 | -1,77 | Transmembrane protein 45B |
| Smp_142970 | -1,77 | -1,24 | -1,76 | palmitoyl protein thioesterase 1 |
| Smp_163630 | -1,44 | -1,12 | -1,69 | developmentally regulated antigen 10 |
| Smp_152940 | -1,61 | -0,97 | -1,60 | otopetrin |
| Smp_126120 | -1,65 | -1,07 | -1,59 | LAMA protein 2 |
| Smp_212450 | -1,30 | -1,20 | -1,51 | placental protein 11 |
| **schistosomula upregulated genes** | | | | |
| **GeneID** | **(Log2FoldChange)** | | | **product_description** |
| | **Cuffdif** | **b-Sleuth** | **EdgeR** | |
| Smp_199840 | 0,73 | 0,48 | 0,89 | nucleolar protein c7b |
| Smp_040990 | 0,98 | 0,57 | 0,81 | Ribonuclease H2 subunit C |
| Smp_149060 | 1,00 | 0,55 | 0,80 | U3 small nucleolar RNA associated protein 22 |
| Smp_190720 | 0,78 | 0,53 | 0,66 | aspartyl tRNA synthetase |
| Smp_140560 | 0,76 | 0,45 | 0,66 | TLC domain containing protein 2 |
| Smp_092390 | 0,62 | 0,44 | 0,61 | N acetylglucosamine kinase |
| Smp_013360 | 0,57 | 0,42 | 0,57 | u1 small nuclear ribonucleoprotein 70 kDa |
| Smp_179010 | 0,64 | 0,40 | 0,56 | exosome complex component RRP42 |
| Smp_124940 | 0,70 | 0,40 | 0,56 | adrenodoxin protein 2C mitochondrial like |
| Smp_208070 | 0,58 | 0,41 | 0,54 | CDC16 cell division cycle 16 |
| Smp_002820 | 0,64 | 0,37 | 0,52 | zinc finger CCCH domain containing protein 4 |
| Smp_170030 | 0,49 | 0,35 | 0,49 | sh3 domain binding glutamic acid rich |
| Smp_022730 | 0,53 | 0,35 | 0,49 | H:ACA ribonucleoprotein complex subunit |
| Smp_042430 | 0,48 | 0,35 | 0,48 | dna replication complex gins protein sld5 |
| Smp_042420 | 0,48 | 0,35 | 0,48 | histone H4 transcription factor |
| Smp_153710 | 0,83 | 0,34 | 0,47 | mutS protein 5 |
| Smp_179160 | 0,53 | 0,41 | 0,47 | nuclear DNA binding protein |
| Smp_071640 | 0,50 | 0,33 | 0,46 | arginine:serine rich splicing factor |
| Smp_126690 | 0,55 | 0,33 | 0,45 | COP9 signalosome complex subunit 6 |
| Smp_058780 | 0,53 | 0,29 | 0,43 | leukocyte receptor cluster |
